# Multicenter self-supervised computational pathology identifies prognostic histomorphological phenotypes in colorectal cancer

**DOI:** 10.64898/2026.07.15.738753

**Authors:** Floor Heilijgers, Hortense Le, Nicolas Coudray, Afreen Karimkhan, Derek Chen, Koen C. M. J. Peeters, Sean Hacking, Wilma E. Mesker, Aristotelis Tsirigos, UNITED collaboration

**Affiliations:** Department of Surgery, Leiden University Medical Center, Leiden, The Netherlands; Division of Precision Medicine, Department of Medicine, Grossman School of Medicine, New York University, New York, New York; Department of Pathology, New York University Grossman School of Medicine, New York, NY, USA; Applied Bioinformatics Laboratories, New York University Grossman School of Medicine, New York, NY, USA

## Abstract

H&E whole-slide images capture prognostic information encoded in tumor morphology and the surrounding microenvironment, but these signals remain difficult to extract and interpret at scale. Here, we developed a self-supervised computational pathology framework to predict disease-free survival in colorectal cancer and link model-derived risk to interpretable histomorphology and spatial tumor biology. Using a multicenter developmental cohort spanning colorectal adenomas and invasive colorectal cancer, we trained HPL-PanColon, a self-supervised representation model, to extract tile-level embeddings and identify recurrent histomorphological phenotype clusters across the adenoma-carcinoma spectrum. Compared with general-purpose pathology foundation models, HPL-PanColon yielded representations with reduced institution- and dataset-specific batch effects. We then applied HPL-PanColon to a global survival cohort of 1,024 colorectal cancer patients in a leave-one-institution-out framework, using tile embeddings to train an attention-based survival model and derive the Colon Histomorphology Prognostic Score (CHiPS). CHiPS stratified patients by disease-free survival and provided complementary prognostic information to a UICC TNM-informed clinicopathological model, increasing the c-index from 0.683 to 0.706. Integrating model attention with phenotype assignments traced CHiPS-associated risk to pathologist-recognizable tissue patterns, with high-risk regions enriched for desmoplastic, stromal, and fibroinflammatory morphologies and low-risk regions reflecting tumor-rich epithelial glandular patterns. Spatial transcriptomic analysis further linked high-risk morphologies to fibroblastic, perivascular, myofibroblastic, and immune-reactive tumor microenvironment programs, while low-risk morphologies mapped to epithelial and tumor-enriched regions. These findings establish a scalable framework for interpretable histology-based prognosis and spatial biological discovery in colorectal cancer.

## Introduction

Colon cancer is among the most prevalent malignancies, ranking as the third-leading cause of cancer-related mortality and accounting for approximately 10% of all cancer diagnoses (1). The TNM (tumor-node-metastasis) classification remains the cornerstone of clinical decision-making and treatment guidelines for colon cancer (2, 3, 4, 5). Despite its limited prognostic capacity, TNM staging remains the basis for assessing survival risk and determining eligibility for adjuvant chemotherapy (ACT) (6). High-risk stage II and III colon cancer patients are routinely treated with surgical resection of the primary tumor followed by ACT, a one-size-fits-all approach that has not been improved in the last 20 years (7). Only about 20% of patients derive meaningful benefit from chemotherapy, while the remaining 80% are either undertreated and could benefit from alternative adjuvant therapy (∼30%), or overtreated and receive ACT despite minimal therapeutic gain (∼50%) thereby being exposed to unnecessary toxicity (3, 4, 8, 9, 10, 11).Currently, we are unable to identify these patients upfront. This emphasizes the clinical need to improve individualized ACT indications through additional biomarkers. Additional pathological parameters that have been implemented in guidelines, such as Tumor Budding and Microsatellite Instability (MSI), mainly focus on the epithelial compartment of the tumor (12, 13, 14).

In recent years, increasing attention has been directed toward the tumor microenvironment (TME) as a critical determinant of tumor behavior (15). In particular, the tumor stroma within the TME has emerged as a key component influencing disease progression, metastatic potential, and therapeutic resistance. A key histological parameter for evaluating this aspect of the TME is the tumor–stroma ratio (TSR), which quantifies the proportion of stroma compared with the tumor epithelial component (16). Accumulating evidence indicates that a high stromal content is strongly associated with adverse clinical outcomes, underscoring the prognostic relevance of stromal architecture (15, 17, 18).

The diagnosis of colon cancer relies on microscopic evaluation of hematoxylin and eosin (H&E)-stained resection specimens by pathologists, making colorectal specimens a substantial component of routine diagnostic workload. Advances in digital pathology, particularly the adoption of whole-slide imaging (WSI), have enabled the large-scale application of artificial intelligence (AI)-based methods to histopathological assessment. AI-based computational tools, applied to digitized tissue sections, assist pathologists in routine tasks such as tumor detection, subtyping, and grading (19). These diagnostic tools, developed for various organs, have already demonstrated benefits, including enhanced prognostic value and reduced workload for pathologists (20, 21, 22). Deep learning (DL) algorithms applied to digitized tissue sections have demonstrated utility in tasks such as tumor detection, subtyping, and grading across multiple organ systems (19). These tools have shown promise in enhancing diagnostic consistency, improving prognostic stratification and reducing pathologist workload (20, 21, 22, 23).

In colorectal cancer, weakly supervised deep learning models have demonstrated that routine H&E histology contains information predictive of molecular pathways and key mutations, including BRAF and KRAS (24). However, many existing approaches are optimized for slide-level prediction and provide limited insight into the recurrent histomorphological phenotypes underlying these associations, constraining biological interpretability and cross-cohort generalizability (24, 25). In contrast, self-supervised learning (SSL) approaches can learn biologically meaningful image representations from unlabeled histology data, providing a scalable framework for phenotype discovery and downstream outcome prediction (26).

We have shown that self-supervised learning enables the extraction of rich histomorphological features from tissue images (27, 28). These features can be organized into morphologic clusters that capture recurrent tissue patterns and may reflect underlying tumor biology. By linking these clusters to clinical outcomes, molecular characteristics, and tumor microenvironment composition, this approach connects image-derived morphology to clinically relevant biology through tissue patterns that can be recognized and interpreted by pathologists (28, 29). By enabling systematic, unbiased characterization of both epithelial and stromal compartments, SSL-based clustering approaches hold substantial potential to refine prognostic stratification and uncover clinically relevant phenotypes in colon cancer.

More broadly, recent pathology foundation models such as Prov-GigaPath, UNI and TITAN illustrate the growing potential of large-scale pretraining to generate reusable histomorphological representations from WSIs (30, 31, 32). These models complement clustering-based SSL approaches by providing generalizable embeddings that can support downstream tasks such as phenotype discovery, survival modeling and tumor microenvironment characterization.

A previous study of Liu et al. (27) employed a SSL-algorithm to derive histomorphological phenotype clusters (HPCs) from unannotated H&E-stained WSIs in colon cancer. These HPCs captured biologically meaningful patterns and were associated with overall survival (OS), outperforming baseline clinical models and providing insights into tumor-stroma composition, immune features and oncogenic pathways. However, disease-free survival (DFS) may more directly reflect tumor biology and recurrence risk compared to OS, as it is less influenced by non-cancer-related mortality and therefore more closely captures disease progression and recurrence. Furthermore, the AVANT trial, analyzed in this cohort, showed treatment heterogeneity, which limited subgroup analysis and generalizability across broader clinical scenarios (27, 28).

Building on these advances, the present study leverages an SSL algorithm (Fig. 1) trained on the multi-institutional, international UNITED cohort (Fig. 1a). The UNITED study validated the prognostic value of the Tumor-Stroma Ratio in stage II and III colon cancer patients (33). This paved the way for TSR’s inclusion in ninth edition of the TNM-classification by The Union for International Cancer Control (UICC) and the American Joint Committee on Cancer (AJCC) (6). The SSL algorithm in this study had the specific objective of predicting survival outcomes from histopathological features, thereby advancing robust, interpretable AI-based prognostication in colon cancer.

**Figure 1.**
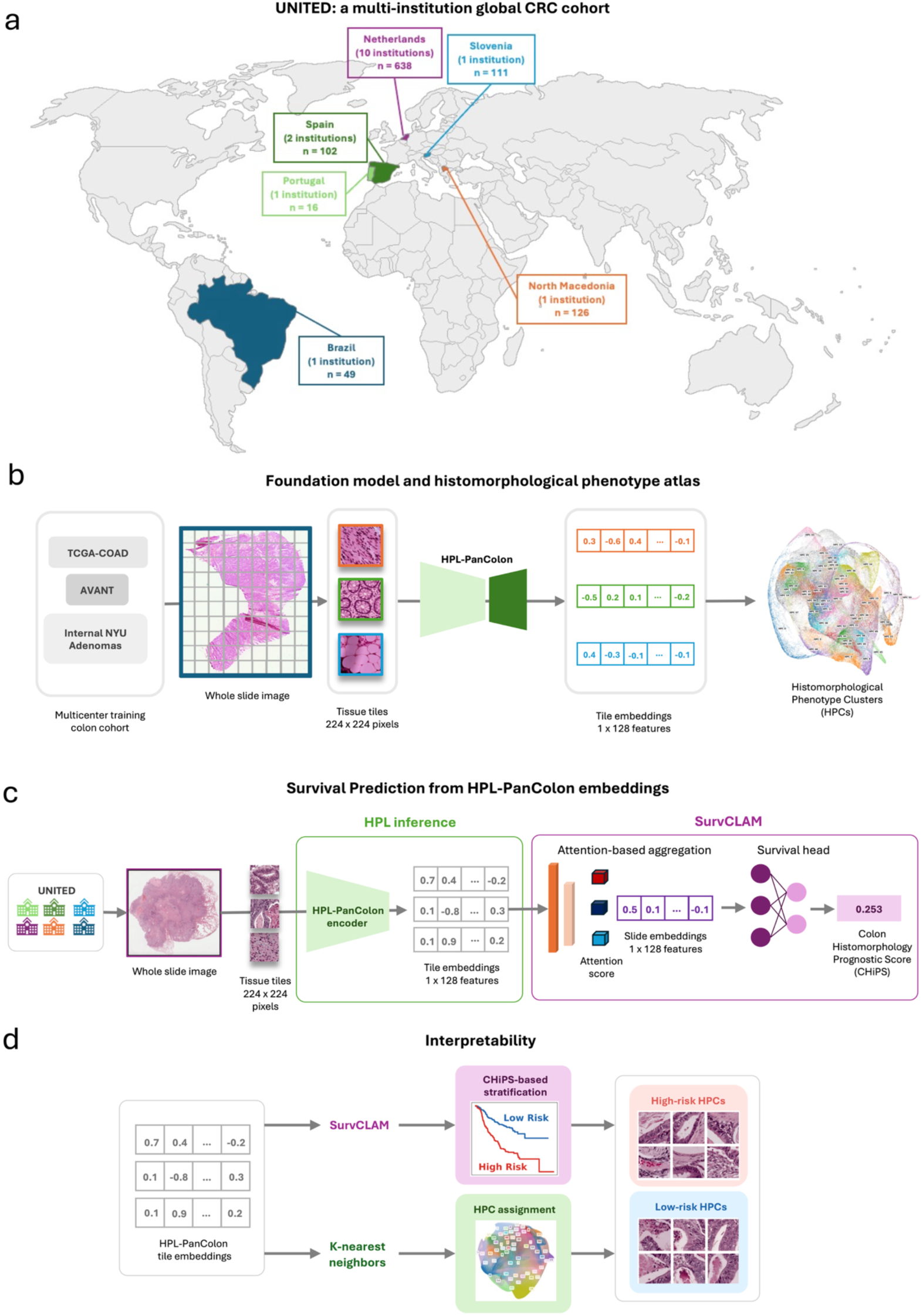
Multi-institutional framework for self-supervised histomorphology learning and survival prediction in colorectal neoplasia. **a,** Geographic distribution of the UNITED cohort, the primary cohort used for survival prediction, comprising 16 institutions across South America and Europe. Numbers indicate the number of contributing institutions and patients in each region. **b,** Development of the HPL-PanColon foundation model. Whole-slide H&E images from TCGA-COAD, the AVANT trial, and internal NYU adenoma cases were tiled and used for self-supervised training to learn a shared histomorphological representation across colorectal neoplasia, from which histomorphological phenotype clusters (HPCs) were defined. **c,** Application of the pretrained HPL-PanColon framework to the UNITED cohort. Tile-level embeddings and HPC assignments were generated from UNITED slides. Embeddings were aggregated with the SurvCLAM survival multiple-instance learning model to derive the patient-level Colon Histomorphology Prognostic Score (CHiPS). **d,** Interpretation of CHiPS-associated histomorphological phenotypes. Tile embeddings from UNITED slides were used in two complementary ways. First, SurvCLAM-derived CHiPS enabled stratification of patients into risk groups. Second, embeddings were mapped to the HPL-PanColon atlas by k-nearest-neighbor assignment to obtain HPC labels. By linking CHiPS-based risk stratification with HPC assignments, we identified the histomorphological phenotypes most strongly associated with high- and low-risk disease, providing biological and morphological interpretability to the survival predictions.

## Results

### HPL-PanColon, a self-supervised colorectal histology model, defines a histomorphological phenotype atlas

We trained HPL-PanColon using colorectal whole-slide images from the TCGA-COAD, AVANT, and ADENOMAS cohorts, which together span colorectal adenomas and invasive colorectal cancer (see Methods). To assess the suitability of its representations for cross-cohort analysis, we compared HPL-PanColon with TITAN (32) and UNI (31). UMAP visualizations showed less apparent cohort-associated structure in the development cohorts and less apparent institution-associated separation in the multicenter UNITED cohort for HPL-PanColon than for TITAN or UNI (Supplementary Figs. 1, 2). We therefore used HPL-PanColon embeddings for downstream phenotype discovery and cross-institutional survival modelling.

Tile-level embeddings from HPL-PanColon were clustered to identify recurrent histomorphological phenotype clusters (HPCs), thereby constructing a colorectal histomorphological phenotype atlas. These HPCs served as interpretable morphological units for downstream survival modelling and biological characterization (Fig. 1b).

### HPL-PanColon supports cross-institutional prediction of disease-free survival in the UNITED cohort

We assessed the prognostic utility of HPL-PanColon embeddings in the multicenter UNITED cohort by applying SurvCLAM to patient-level bags of slide embeddings in a leave-one-institution-out framework across 16 institutions to predict DFS (Fig. 1c). This approach yielded a continuous patient-level risk score, CHiPS, from pooled out-of-fold predictions.

CHiPS stratified patients into distinct prognostic groups. When pooled out-of-fold CHiPS values were divided into tertiles, Kaplan–Meier analysis demonstrated clear separation of low-, intermediate- and high-risk groups, with progressively worse DFS across increasing CHiPS strata (Fig. 2a). These findings indicate that histomorphological signals captured by HPL-PanColon and aggregated by SurvCLAM are sufficient to identify patients with different clinical outcomes.

**Figure 2.**
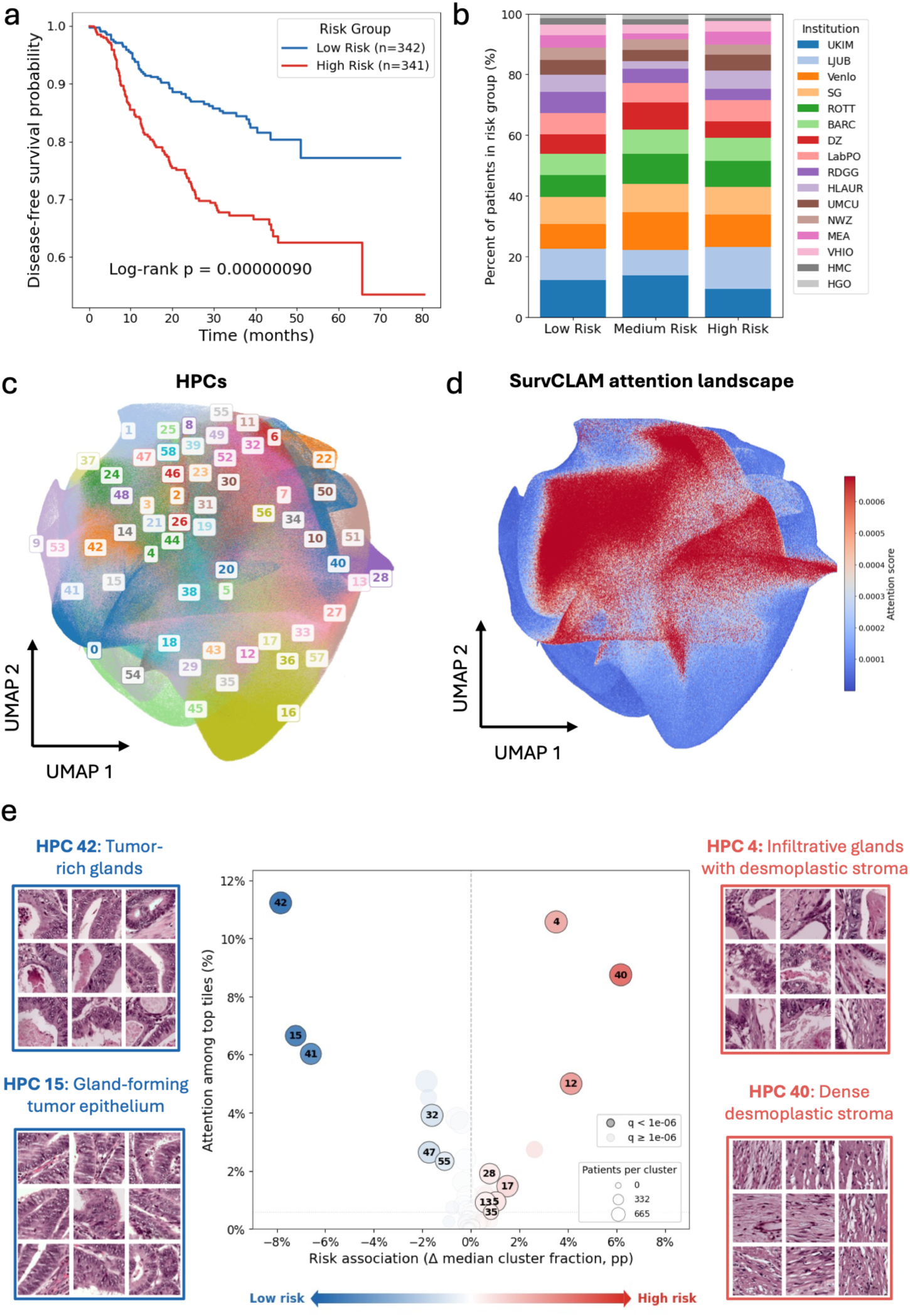
Histomorphological phenotypes associated with survival risk and model attention in the UNITED cohort. **a,** Kaplan–Meier curves for patients stratified into low-, and high-risk groups according to the SurvCLAM-derived Colon Histomorphology Prognostic Score (CHiPS), defined as the lowest and highest score tertiles, demonstrating significant separation of disease-free survival across risk strata. **b,** Institutional composition of CHiPS risk groups in the UNITED cohort. Bars show the percentage of patients contributed by each institution within each risk stratum. **c,** Projection of UNITED tile embeddings into the HPL-PanColon histomorphological phenotype atlas. Colors denote histomorphological phenotype clusters (HPCs), and numbers indicate individual HPCs. **d,** SurvCLAM attention mapped onto the same latent space, showing that highly attended tiles localize to discrete regions of morphology space rather than being uniformly distributed across the atlas. Attention weights are clipped at the 1st and 99th percentiles across all tiles cohort-wide for visualization. **e,** Association of HPCs with survival risk and model attention. Each point denotes one HPC, plotted according to its differential abundance between high- and low-risk CHiPS tertiles, defined as the difference in median cluster fraction (in percentage points) between patients in the highest- and lowest-risk groups, and its representation among the top 25% highest-attention tiles. Positive x-axis values indicate enrichment in high-risk patients, whereas negative values indicate enrichment in low-risk patients. Point size corresponds to the number of patients represented by each cluster, and color indicates statistical significance. Only clusters with *q* < 1 × 10⁻⁵ are labeled for readability. Representative tiles illustrate low-risk-associated epithelial and gland-forming phenotypes, including tumor-rich glands and gland-forming tumor epithelium, and high-risk-associated stromal phenotypes, including infiltrating glands with desmoplastic stroma and dense desmoplastic stroma.

We next examined whether this separation might reflect institution-level imbalance. The institutional composition of the low-, intermediate- and high-risk groups was broadly similar (Fig. 2b), arguing against center-specific sampling effects as the principal driver of prognostic stratification. Together, these results support that CHiPS captures a survival-associated signal that generalizes across institutions.

### CHiPS is associated with distinct histomorphological phenotypes prioritized by the survival model

To define the tissue patterns underlying CHiPS, we related tile-level SurvCLAM attention scores to the HPL-PanColon HPCs (Fig. 2c). Projection of UNITED tiles into the phenotype atlas showed that highly attended tiles concentrated in discrete regions of morphology space rather than being diffusely distributed (Fig. 2d), indicating that the model prioritized a restricted set of recurrent tissue phenotypes. To support biological interpretation of the unsupervised phenotype clusters, representative tiles from all 59 HPCs were reviewed by a pathologist and assigned concise morphological annotations, providing a reference atlas for downstream analyses (Supplementary Fig. 3).

We then quantified, for each HPC, its association with CHiPS risk strata and its representation among highly attended tiles (Fig. 2e). Multiple HPCs showed significant associations with risk after adjustment, supporting a broader prognostic histomorphology landscape rather than dependence on a single phenotype. Significant low-risk-associated HPCs included HPC14, HPC15, HPC18, HPC24, HPC25, HPC32, HPC37, HPC41, HPC42, HPC44, HPC47, HPC48, and HPC55. Significant high-risk-associated HPCs included HPC4, HPC5, HPC11, HPC12, HPC13, HPC16, HPC17, HPC28, HPC33, HPC34, HPC35, HPC38, HPC40, and HPC51. Among the most significant and interpretable clusters, low-risk-associated HPC42, HPC15, and HPC41 represented epithelial and gland-forming phenotypes, whereas high-risk-associated HPC4, HPC40, and HPC12 represented stromal, fibromuscular, and desmoplastic phenotypes.

Pathologist review of representative tiles supported these associations (Fig. 2e, Supplementary Fig. 3). Low-risk-associated clusters were characterized by tumor-rich or gland-forming epithelial architecture with minimal intervening stroma, including tumor-rich neoplastic glands with luminal necrosis in HPC42, gland-forming tumor epithelium with minimal desmoplastic stroma in HPC15, and tubular adenoma-like adenomatous epithelium in HPC41. In contrast, high-risk-associated clusters were characterized by stromal and desmoplastic tissue patterns, including infiltrating malignant glands with a prominent desmoplastic stromal background in HPC4, dense hypocellular fibrotic/desmoplastic stroma with limited visible tumor epithelium in HPC40, and dense fibromuscular or fibroblastic stromal tissue in HPC12. Together, these findings suggest that CHiPS reflects an atlas spanning epithelial, and gland-forming phenotypes associated with favorable outcome and stromal, fibromuscular, and desmoplastic phenotypes associated with unfavorable outcome.

### Representative cases reveal contrasting spatial organization of risk-associated morphology

To visualize how CHiPS-associated phenotypes contributed to patient-level risk, we examined representative high- and low-CHiPS cases using spatial overlays of HPC assignments and SurvCLAM attention maps (Fig. 3). High-CHiPS cases, marked by positive CHiPS values, showed broader spatial involvement of high-risk-associated stromal and desmoplastic phenotypes, with model attention concentrated in corresponding regions of the tissue. By contrast, low-CHiPS cases, marked by negative CHiPS values, showed limited attention to high-risk-associated stromal/desmoplastic phenotypes. Notably, low CHiPS did not always require visible enrichment of the low-risk-associated HPCs highlighted in the atlas; in some cases, low predicted risk appeared to reflect the relative absence of high-risk morphology rather than dominance of a specific favorable phenotype.

**Figure 3.**
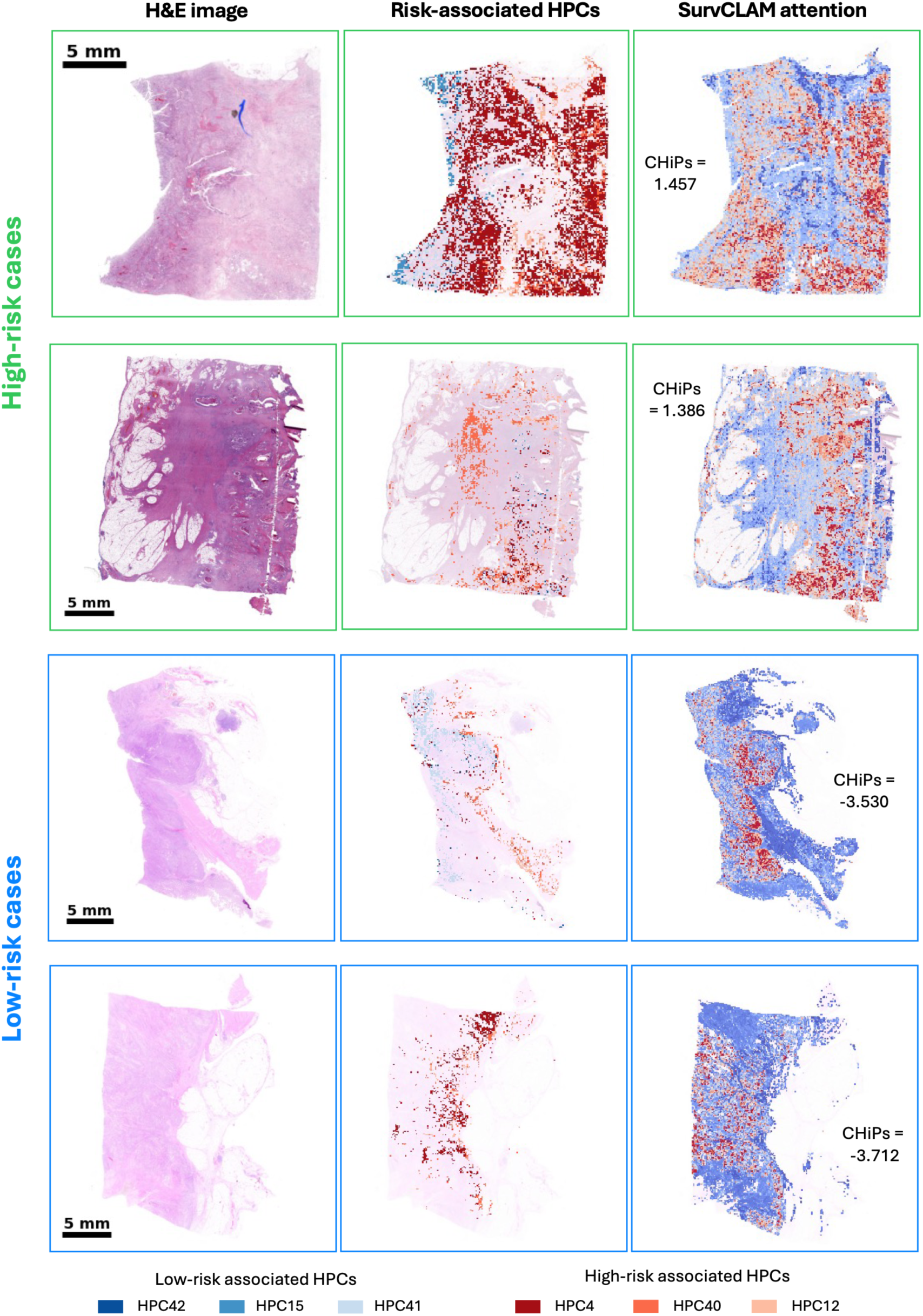
Spatial visualization of CHiPS-associated histomorphological patterns in representative colorectal cancer cases. Representative whole-slide H&E images are shown for high- and low-CHiPS cases together with spatial overlays of selected survival-associated histomorphological phenotype clusters and SurvCLAM attention maps. The overlays show the top selected HPCs per risk group and are intended as illustrative examples rather than exhaustive maps of all phenotypes contributing to CHiPS. High-CHiPS cases show broader spatial representation of high-risk-associated stromal and desmoplastic phenotypes, with model attention concentrated in corresponding tissue regions. In contrast, low-CHiPS cases show limited attention to high-risk-associated stromal/desmoplastic phenotypes; in some cases, low CHiPS appears to reflect the relative absence of high-risk morphology rather than strong enrichment of the selected low-risk HPCs highlighted in the phenotype atlas. These examples illustrate how CHiPS integrates spatially distributed histomorphological patterns across whole-slide images to generate patient-level prognostic risk scores. CHiPS, Colon Histomorphology Prognostic Score; HPC, histomorphological phenotype cluster.

These case-level visualizations support the interpretation that CHiPS is not driven solely by isolated local features or by the presence of a single protective HPC. Instead, CHiPS reflects how SurvCLAM integrates the broader attention-weighted spatial organization of histomorphological states across whole-slide images, including both the presence of high-risk stromal/desmoplastic tissue patterns and their relative absence.

### CHiPS captures prognostic information beyond standard clinicopathological variables

We next examined the relationship between CHiPS and standard clinicopathological variables (Fig. 4, Supplementary Fig. 4). CHiPS was not significantly associated with sex, but differed significantly across pathological T category, pathological N category, and tumor stroma ratio, with higher CHiPS values observed in groups with more adverse pathological features (Fig. 4a). CHiPS also showed a weak negative correlation with age, while summary association analyses confirmed stronger positive associations with pathological T category, pathological N category, and tumor stroma ratio than with sex or age (Supplementary Fig. 4). These findings indicate that CHiPS partially reflects established prognostic morphology and disease extent, while also motivating formal evaluation of whether it provides information beyond conventional clinicopathological variables.

**Figure 4.**
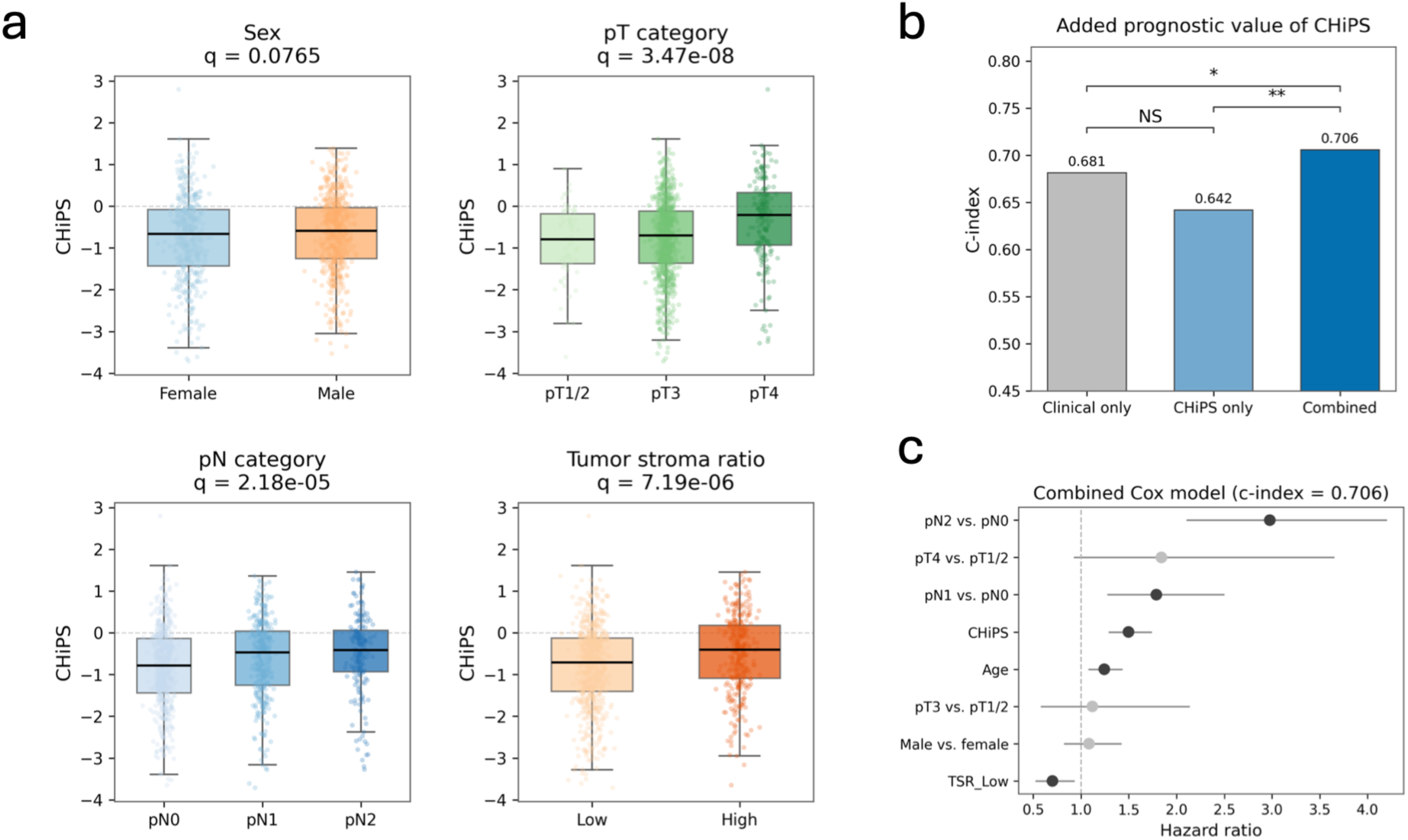
Association of CHiPS with clinicopathologic variables and prognostic performance. **a**, Distribution of CHiPS across standard clinicopathological variables, including sex, pathological T category, pathological N category, and tumor stroma ratio. CHiPS was not significantly associated with sex, but differed significantly across pathological T category, pathological N category, and tumor stroma ratio, indicating partial correspondence with established adverse pathological features. **b**, Comparison of concordance indices for models based on clinicopathological variables alone, CHiPS alone, and the combined model. The combined model achieved the highest concordance index, supporting the added prognostic value of CHiPS beyond standard clinicopathological parameters. Statistical significance for c-index comparisons was assessed by bootstrap resampling of patients across 1,000 iterations; NS, not significant; *P < 0.05; **P < 0.01; ***P < 0.001. **c**, Multivariable Cox proportional hazards model including CHiPS and clinicopathological covariates. CHiPS remained an independent prognostic variable when evaluated alongside conventional clinical and pathological features. CHiPS, Colon Histomorphology Prognostic Score.

We therefore compared prognostic models based on clinicopathological variables alone, CHiPS alone, or both in combination. A model based on clinicopathological variables alone achieved a concordance index of 0.683, whereas CHiPS alone achieved a concordance index of 0.642 (Fig. 4b). Combining CHiPS with clinicopathological variables increased performance to 0.706, outperforming either clinicopathological variables or CHiPS alone. In the multivariable Cox model, CHiPS remained prognostic when evaluated alongside pathological T category, pathological N category, age, sex, and tumor stroma ratio (Fig. 4c).

These results indicate that CHiPS captures histomorphological information that overlaps partially with established adverse clinicopathological features but is not reducible to them. Its improved performance in the combined model supports CHiPS as a complementary prognostic biomarker that can augment, rather than replace, standard clinicopathological risk assessment.

### Spatial transcriptomic profiling links risk-associated phenotypes to distinct cellular and molecular programs

To define the biological basis of the prognostic phenotypes, we mapped HPL-PanColon HPCs onto a public Visium HD colorectal cancer dataset (Fig. 5a). This analysis focused on spatially represented survival-associated HPCs, including low-risk-associated HPC42 and HPC41 and high-risk-associated HPC4, HPC40, and HPC12. Although HPC15 was also associated with lower risk in the UNITED analysis, it was represented by only a single tile across the Visium regions analyzed and was therefore excluded from the spatial transcriptomics analysis.

**Figure 5.**
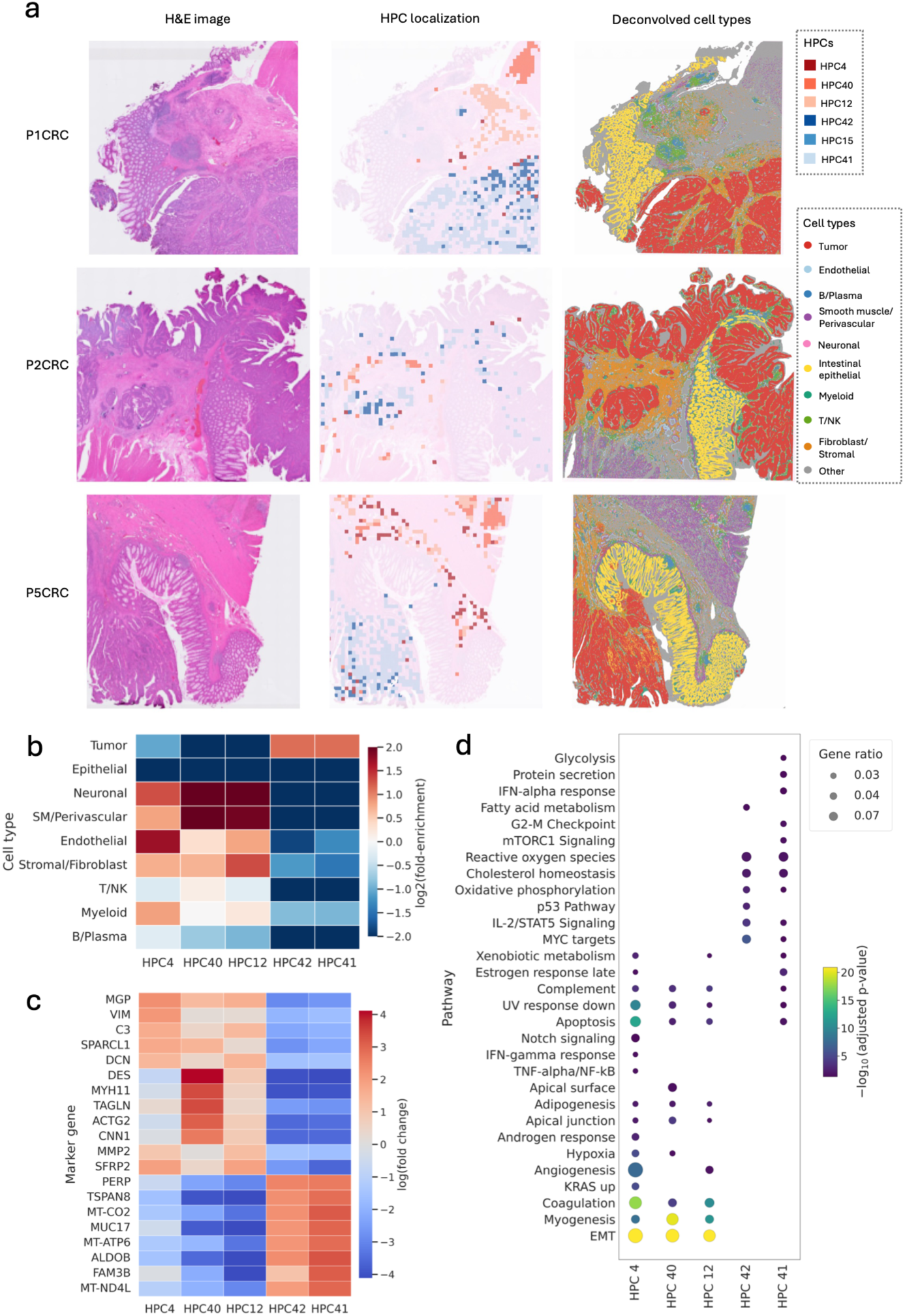
Spatial transcriptomic characterization of survival-associated histomorphological phenotypes in colorectal cancer. **a,** Representative Visium HD spatial transcriptomic sections showing survival-associated HPL-PanColon histomorphological phenotype clusters overlaid on matched H&E images and broad cell-type maps. Broad cell-type maps were derived from published deconvolution results from the spatial transcriptomics study. Low-risk-associated HPC42 and HPC41 localized to tumor-rich glandular and epithelial regions, whereas high-risk-associated HPC4, HPC40, and HPC12 localized to stromal, desmoplastic, fibromuscular, or mesenchymal tissue compartments. **b,** Broad cell-type enrichment across survival-associated HPCs, shown as log2 fold-enrichment relative to the global cell-type distribution. HPC42 and HPC41 were enriched for tumor and epithelial compartments and relatively depleted of stromal-associated cell states. In contrast, HPC4, HPC40, and HPC12 showed enrichment for stromal, fibroblast, smooth muscle/perivascular, or immune-associated compartments. **c,** Marker gene expression across selected survival-associated HPCs. The top five upregulated marker genes were selected for each HPC and then visualized across all selected HPCs as log fold change values, enabling comparison of marker specificity across phenotypes. Low-risk-associated HPC42 and HPC41 displayed epithelial- and glandular-associated marker enrichment, whereas high-risk-associated HPC4, HPC40, and HPC12 showed stromal, mesenchymal, fibroblastic, or contractile marker programs. **d,** Hallmark pathway over-representation analysis across selected survival-associated HPCs using Hallmark gene sets. Only significantly enriched pathways are shown, using an adjusted *P* < 0.05 threshold. Dot size indicates gene ratio, and color indicates enrichment significance as −log10 adjusted *P* value. Low-risk-associated HPC42 was enriched for epithelial, metabolic, and proliferative programs, whereas high-risk-associated stromal/desmoplastic HPCs showed enrichment for Hallmark pathways related to epithelial–mesenchymal transition, myogenesis, coagulation, hypoxia, extracellular matrix organization, and inflammatory signaling. CHiPS, Colon Histomorphology Prognostic Score; HPC, histomorphological phenotype cluster.

The spatial localization of these HPCs supported the histological interpretations from the UNITED cohort. Low-risk-associated HPC42 and HPC41 localized predominantly to tumor-rich glandular and epithelial regions. In contrast, high-risk-associated HPC4, HPC40, and HPC12 localized to stromal, desmoplastic, fibromuscular, and mesenchymal tissue compartments. Broad cell-type enrichment analysis further supported these spatial patterns (Fig. 5b). HPC42 and HPC41 were enriched for tumor and epithelial compartments and relatively depleted of stromal-associated cell states, whereas HPC4, HPC40, and HPC12 showed enrichment for stromal, fibroblast, smooth muscle/perivascular, or immune-associated compartments. Extended analysis across the broader set of CHiPS-associated HPCs showed similar cellular compartment patterns, with low-risk-associated clusters enriched for tumor or epithelial cell states and high-risk-associated clusters enriched for stromal/fibroblast, smooth muscle/perivascular, endothelial, myeloid, or T/NK compartments (Supplementary Fig. 5).

Gene expression and over-representation (ORA) analyses further distinguished these phenotypes at the molecular level (Fig. 5c,d). Low-risk-associated HPC42 and HPC41 showed epithelial and glandular marker programs, whereas high-risk-associated HPC4, HPC40, and HPC12 showed stromal, mesenchymal, fibroblastic, and contractile marker programs. Marker gene analysis across the broader set of CHiPS-associated HPCs further supported this epithelial–stromal distinction, with epithelial/glandular clusters enriched for tumor-associated markers and stromal/desmoplastic clusters enriched for fibroblastic, extracellular matrix, contractile, and immune-associated markers (Supplementary Fig. 5). Consistent with this pattern, pathway enrichment analysis linked low-risk-associated phenotypes to epithelial, metabolic, and proliferative programs, while high-risk-associated stromal/desmoplastic phenotypes were enriched for pathways related to epithelial–mesenchymal transition, myogenesis, coagulation, hypoxia, extracellular matrix organization, and inflammatory signaling. Extended Hallmark ORA across CHiPS-associated HPCs revealed recurrent enrichment of epithelial, metabolic, and proliferative programs among low-risk-associated phenotypes and stromal, mesenchymal, inflammatory, hypoxic, and coagulation-related programs among high-risk-associated phenotypes (Supplementary Fig. 6).

These findings indicate that the prognostic signal captured by CHiPS is anchored in biologically distinct tissue states. Low-risk CHiPS-associated morphologies correspond to tumor-rich epithelial and glandular programs, whereas high-risk CHiPS-associated morphologies correspond to stromal, desmoplastic, fibromuscular, and inflammatory programs. This spatial transcriptomic analysis therefore provides independent biological support for the histomorphological risk patterns identified by HPL-PanColon and prioritized by SurvCLAM.

## Discussion

### Validation survival prediction

In this study, CHiPS enabled robust cross-institutional stratification of disease-free survival and provided prognostic information beyond standard clinicopathological variables. Risk groups were driven by distinct histomorphological patterns, with favorable outcomes associated with epithelial and gland-forming phenotypes and poorer outcomes linked to stromal and desmoplastic states. Rather than relying on isolated features, CHiPS captured the spatial organization of tissue phenotypes across whole slides. Integration with spatial transcriptomics further demonstrated that these morphologies correspond to distinct molecular programs, supporting the biological interpretability of CHiPS and highlighting its potential value for improved risk assessment in colon cancer.

### Correlation TME/TSR

As was shown in Liu et al. (27), this study again emphasized the importance of the TME, particularly tumor stroma, in shaping patient outcomes. Tumor stroma, composed of extracellular matrix, vasculature, immune cells and cancer-associated fibroblasts, forms a dynamic and complex interaction with tumor epithelial cells (15, 34). High-risk CHiPS-associated phenotypes were characterized by stromal and desmoplastic morphologies, supporting the established prognostic role of stromal abundance, architecture, and immune infiltration (17, 33, 35, 36). Spatial transcriptomic analyses demonstrated that CHiPS captures higher-order tissue organization rather than isolated local features, indicating that its prognostic signal is rooted in biologically distinct epithelial and stromal tissue states. Interestingly, some low-CHiPS cases showed limited representation of the selected low-risk-associated HPCs highlighted in the spatial overlays, suggesting that favorable CHiPS predictions may arise not only from enrichment of the most prominent epithelial and glandular phenotypes but also from the relative absence of high-risk stromal/desmoplastic programs. Because the overlays visualize only the top selected HPCs per risk group, these examples should be interpreted as illustrative rather than exhaustive representations of the morphologies contributing to CHiPS.

From a clinical perspective, these findings are particularly relevant when considering established stromal biomarkers such as the tumor stroma ratio (TSR). Although CHiPS was strongly associated with TSR, it extends beyond a simple estimate of stromal content by incorporating detailed histomorphological information, including stromal heterogeneity and the spatial relationships between epithelial, stromal, and immune compartments. This likely explains the improved prognostic performance observed when CHiPS was combined with standard clinicopathological variables.

Moreover, spatial transcriptomic analyses revealed that adverse morphologies corresponded to complex reactive microenvironmental programs involving fibroblast, vascular, and immune components, whereas favorable epithelial phenotypes represented multiple distinct tumor-rich states. Together, these findings suggest that CHiPS captures multiple favorable and unfavorable tissue states rather than a single protective or adverse morphology.

### Clinical utility

The present findings highlight the potential clinical utility of CHiPS and its underlying atlas of histomorphological phenotypes. By providing prognostic information beyond established clinicopathological factors, CHiPS may improve risk stratification, particularly in settings where current markers offer limited guidance, such as stage II colon cancer. Current treatment of stage II colon cancer relies primarily on surgical resection, with ACT reserved for patients considered at high risk of recurrence (2, 3, 4, 5). High-risk features typically include pT4 tumors, inadequate lymph node sampling, poor differentiation, high tumor budding, lympho-angioinvasion, and clinical factors such as obstruction or perforation (4). Despite these criteria, accurately identifying patients who will benefit from ACT remains challenging, resulting in both overtreatment and undertreatment of stage II patients (3, 4, 8, 9, 10, 11). Identification of high-risk stromal phenotypes could support more personalized decisions regarding adjuvant therapy and follow-up, while low-risk glandular phenotypes may help avoid overtreatment.

Importantly, the biologically interpretable nature of the identified HPCs facilitates clinical adoption by linking risk predictions to recognizable histomorphological patterns rather than functioning as a “black box”. Integrated alongside routine pathology assessment (Fig. 6), CHiPS could provide complementary prognostic information while reducing the burden of complex pattern recognition.

**Figure 6.**
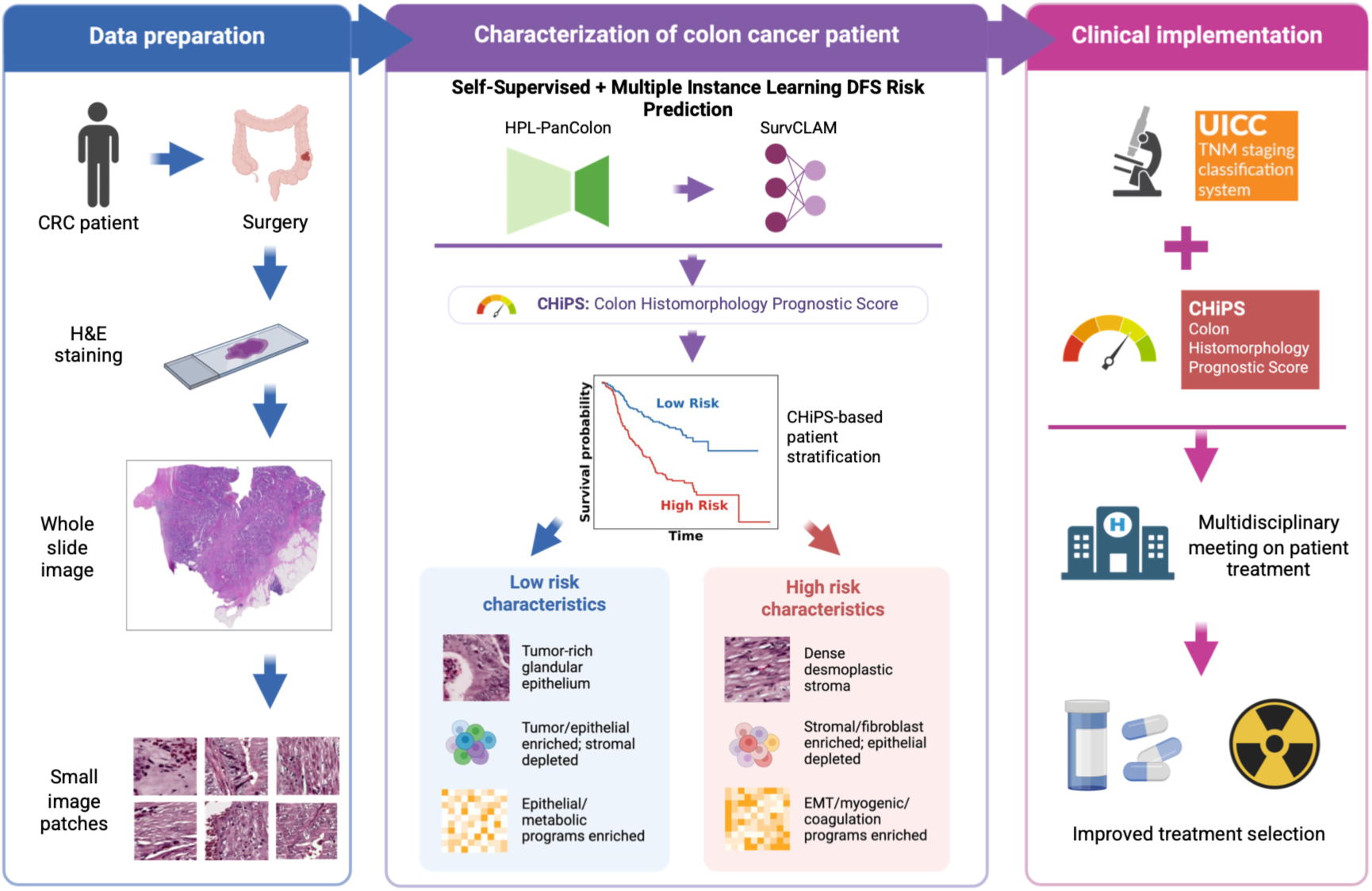
Clinical implementation of histomorphology-based survival prediction in colorectal cancer. **Left,** routine pathology processing from surgical resection to H&E staining, whole-slide imaging, and tile extraction. **Middle,** tile-level features are extracted with the HPL-PanColon foundation model and aggregated by the SurvCLAM survival multiple-instance learning framework to derive the Colon Histomorphology Prognostic Score (CHiPS), enabling risk stratification and interpretation through associated histomorphological phenotypes. **Right,** CHiPS is integrated with standard clinicopathological assessment to support multidisciplinary treatment planning and patient management. Figure created with BioRender.com. CRC: colorectal cancer; DFS: disease-free survival; EMT: Epithelial-Mesenchymal Transition; ECM: Extracellular Matrix

Nevertheless, the current findings are primarily prognostic, and CHiPS cannot yet be used to guide treatment decisions directly. Further validation in prospective and treatment-stratified cohorts is required to establish its predictive value and clinical utility.

### Limitations

Several limitations should be considered when interpreting these findings. First, although the multicenter, leave-one-institution-out design supports generalizability, the analysis remains retrospective and is therefore susceptible to residual confounding and selection biases. Second, while CHiPS provided complementary prognostic information when combined with standard clinicopathological variables, the incremental performance gain over clinicopathological variables alone was modest and requires validation in larger prospective cohorts. In addition, performance may be affected by interinstitutional variability in tissue processing, staining, and slide digitization.

The spatial transcriptomic analyses were based on a limited number of Visium HD sections and relied on deconvolution-based annotations, which may not fully capture the molecular heterogeneity and cellular complexity of colorectal cancer. As a result, the molecular interpretation should be viewed as supportive rather than exhaustive. Finally, the biological mechanisms underlying the identified tissue states remain incompletely understood, and further prospective validation and workflow standardization will be required before clinical implementation.

### Future perspectives

Future research should focus on gaining deeper insight into the clinically relevant, pixel-level features that drive AI-generated HPC formation. Improved interpretability of these features will be essential to strengthen trust and facilitate integration into routine pathology workflows. In addition, the development of AI tools capable of analyzing entire tumor resection specimens, rather than one WSI per patient, could enable more comprehensive tumor characterization and yield robust, patient-level risk scores. Such risk stratification tools hold the potential to guide individualized treatment decisions, ultimately helping to reduce both overtreatment and undertreatment. To achieve true clinical impact, future studies should prioritize prospective validation and assess the predictive value of HPC-based models for treatment response across different therapeutic modalities.

## Conclusion

In conclusion, we demonstrate that CHiPS enables robust prediction of DFS and stratifies stage II-III colorectal cancer patients into clinically meaningful risk groups. This prognostic signal is rooted in a reproducible and biologically interpretable spectrum of HPCs, ranging from tumor-rich glandular states to stromal and desmoplastic tissue programs with distinct spatial organization and molecular characteristics. The results support the impact of the TME and specifically the stromal features. Importantly, CHiPS captures prognostic information beyond standard clinicopathological variables. Together, these findings underscore the potential of integrating deep learning–based histomorphological profiling into clinical practice to improve individualized risk assessment and support treatment decision-making in colorectal cancer.

## Methods

### Ethics statement

Our research complies with all relevant ethical regulations. The study protocol was approved by Applied Bioinformatics Laboratories of New York University Grossman School of Medicine and the Department of Surgery of Leiden University Medical Center. The analyses were performed using anonymized material, and all patients approved for the study including AI and automated digital analysis (33). Data from TCGA-COAD was open-accessed, ensuring patient anonymity without risk of patient identification. All institutions contributing annotated biospecimens provided documentation to the TCGA, and have obtained ethical approvals to use the sample and data according to the human subjects protection and data access policies in TCGA program. Archival material derived from the AVANT-trial (BO17920) was performed in accordance with the declaration of Helsinki (37, 38). Protocol approval was obtained from the local medical ethics review committees or institutional review boards at participating sites.

### Patient population

This study leveraged multiple colorectal histopathology cohorts spanning adenomas and adenocarcinomas. HPL-PanColon (Fig. 1b), a self-supervised model for learning histomorphological representations from H&E whole-slide images (28), was trained using TCGA-COAD, the Bevacizumab-Avastin® adjuVANT (AVANT) trial, and an internal ADENOMAS cohort from NYU, and tested using the UNITED cohort.

The final TCGA-COAD dataset included 435 WSIs from 428 patients with a diagnosed pathological TNM-stage I-IV colon carcinoma (27, 39). The AVANT cohort, comprising 1213 colon cancer patients with available diagnostic H&E WSIs (one WSI per patient), was used, due to the previously studied potential correlation of the stromal compartment (eg. TSR) and patient prognosis (35, 37, 38). For a detailed overview of the AVANT trial and patient characteristics, see De Gramont et al. (38). Given the unique treatment regimen of bevacizumab, we only utilized the control group that received FOLFOX-4 (without bevacizumab), ensuring treatment consistency between cohorts. The details on above mentioned cohorts are described in our previous study (27). The ADENOMAS cohort comprised an unpublished internal dataset of 832 whole-slide images from 309 patients, including 122 patients who progressed and 187 patients who did not progress. Downstream survival prediction was performed in the UNITED cohort (Fig. 1a). The original cohort comprised 1,388 patients from 27 institutions across 12 countries. Due to limited slide availability and slide corruption, 1,058 slides from 1,028 patients representing 16 institutions were ultimately included in this study. For biological interpretation of the learned phenotypes, we additionally analyzed a public Visium HD spatial transcriptomic colorectal cancer dataset from 3 patients with matched H&E images and spatially resolved gene expression profiles (40).The eligible UNITED cohort consisted of 1028 neoadjuvant naïve, stage II and III colon cancer patients. All patients had received a curative resection (R0) of their primary tumor, and 48.4% (n = 494 patients) had received ACT. The median age was 68 years, and 569 patients (55.8%) were of male sex. A total of 519 patients (50.9%) had stage II colon cancer. Additional baseline characteristics are shown in Table 1. Detailed information regarding the UNITED study can be found elsewhere (33).

**Table 1.**
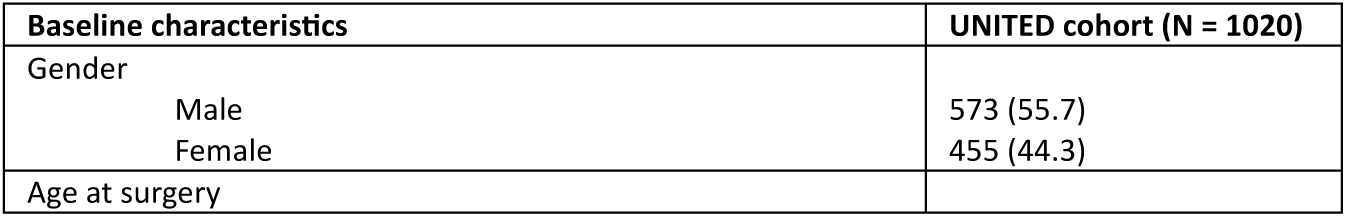
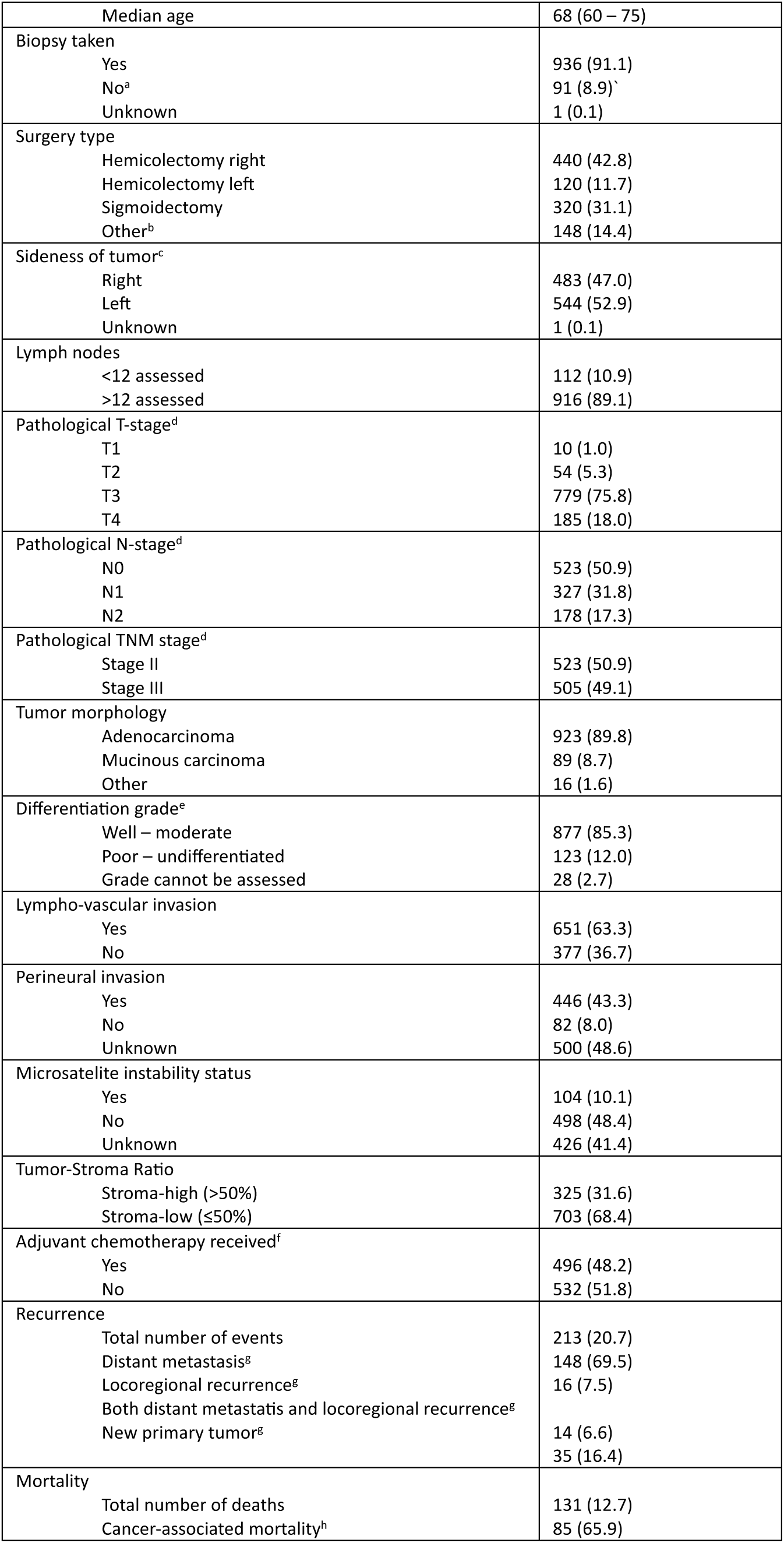
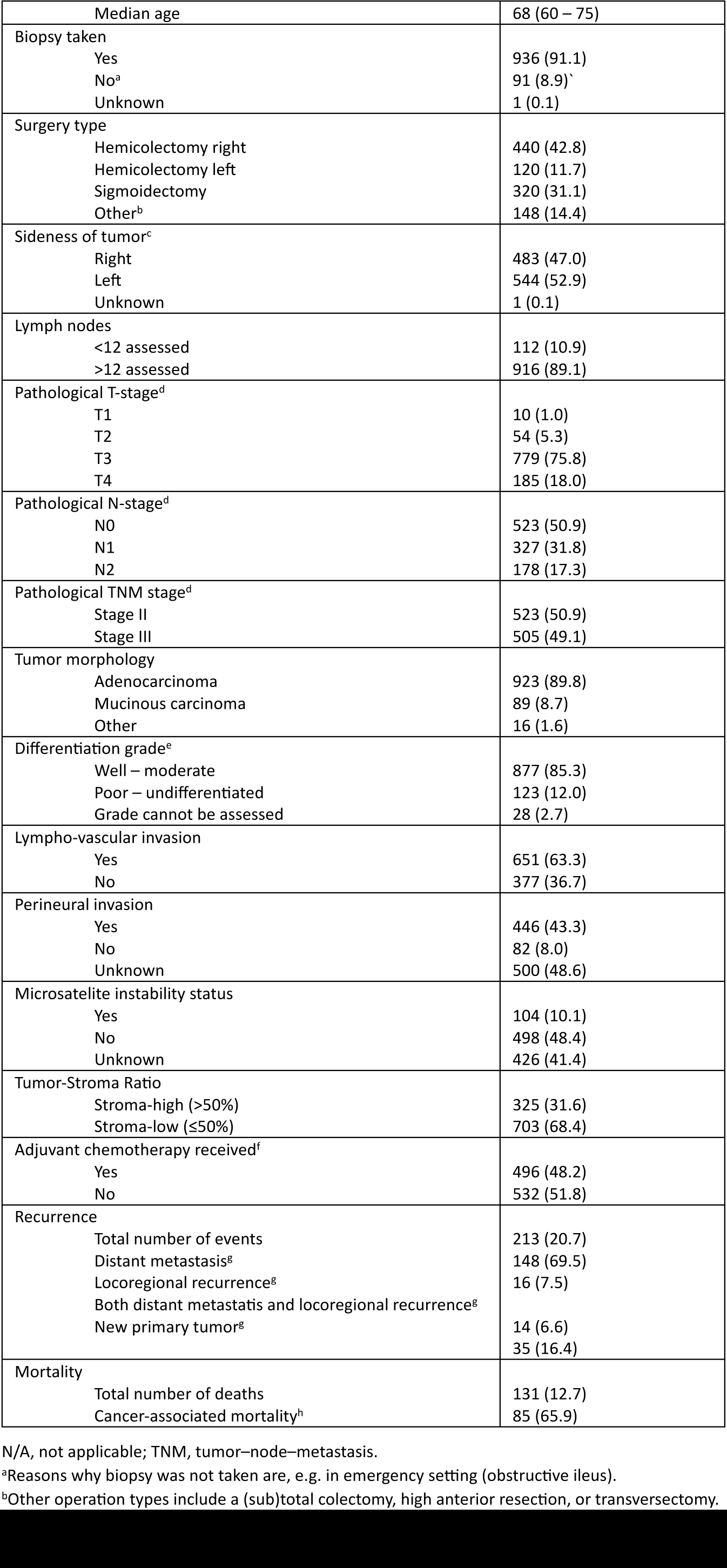
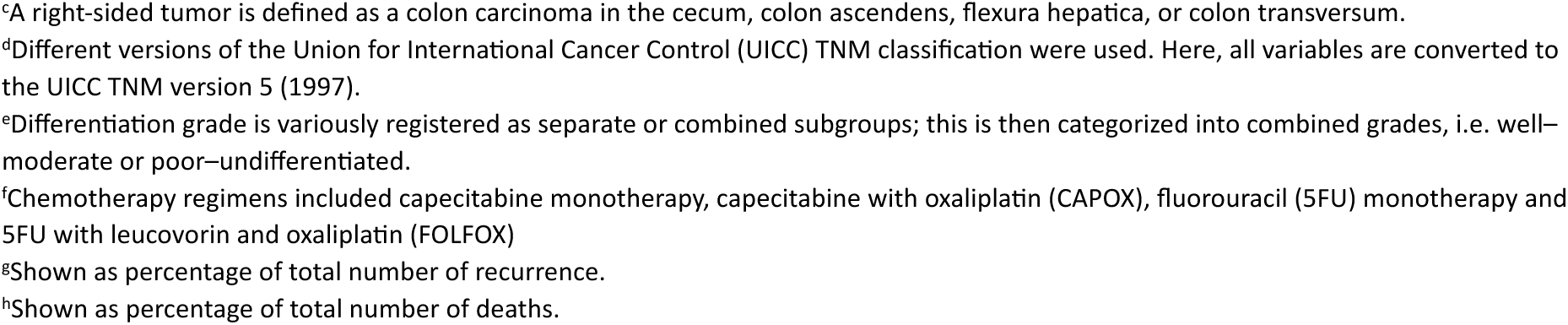
Baseline characteristics of patients in the eligible UNITED cohort. All variables are given as absolute numbers with associated percentages or medians with interquartile ranges. Sum of percentages can be less or more than 100 due to rounding.

| Baseline characteristics | UNITED cohort (N = 1020) |
| --- | --- |
| Gender |  |
| Male | 573 (55.7) |
| Female | 455 (44.3) |
| Age at surgery |  |
| Median age | 68 (60 – 75) |
| Biopsy taken |  |
| Yes | 936 (91.1) |
| No<sup>a</sup> | 91 (8.9) |
| Unknown | 1 (0.1) |
| Surgery type |  |
| Hemicolectomy right | 440 (42.8) |
| Hemicolectomy left | 120 (11.7) |
| Sigmoidectomy | 320 (31.1) |
| Other<sup>b</sup> | 148 (14.4) |
| Sideness of tumor<sup>c</sup> |  |
| Right | 483 (47.0) |
| Left | 544 (52.9) |
| Unknown | 1 (0.1) |
| Lymph nodes |  |
| <12 assessed | 112 (10.9) |
| >12 assessed | 916 (89.1) |
| Pathological T-stage<sup>d</sup> |  |
| T1 | 10 (1.0) |
| T2 | 54 (5.3) |
| T3 | 779 (75.8) |
| T4 | 185 (18.0) |
| Pathological N-stage<sup>d</sup> |  |
| N0 | 523 (50.9) |
| N1 | 327 (31.8) |
| N2 | 178 (17.3) |
| Pathological TNM stage<sup>d</sup> |  |
| Stage II | 523 (50.9) |
| Stage III | 505 (49.1) |
| Tumor morphology |  |
| Adenocarcinoma | 923 (89.8) |
| Mucinous carcinoma | 89 (8.7) |
| Other | 16 (1.6) |
| Differentiation grade<sup>e</sup> |  |
| Well – moderate | 877 (85.3) |
| Poor – undifferentiated | 123 (12.0) |
| Grade cannot be assessed | 28 (2.7) |
| Lympho-vascular invasion |  |
| Yes | 651 (63.3) |
| No | 377 (36.7) |
| Perineural invasion |  |
| Yes | 446 (43.3) |
| No | 82 (8.0) |
| Unknown | 500 (48.6) |
| Microsatellite instability status |  |
| Yes | 104 (10.1) |
| No | 498 (48.4) |
| Unknown | 426 (41.4) |
| Tumor-Stroma Ratio |  |
| Stroma-high (>50%) | 325 (31.6) |
| Stroma-low (≤50%) | 703 (68.4) |
| Adjuvant chemotherapy received<sup>f</sup> |  |
| Yes | 496 (48.2) |
| No | 532 (51.8) |
| Recurrence |  |
| Total number of events | 213 (20.7) |
| Distant metastasis<sup>g</sup> | 148 (69.5) |
| Locoregional recurrence<sup>g</sup> | 16 (7.5) |
| Both distant metastasis and locoregional recurrence<sup>g</sup> | 14 (6.6) |
| New primary tumor<sup>g</sup> | 35 (16.4) |
| Mortality |  |
| Total number of deaths | 131 (12.7) |
| Cancer-associated mortality<sup>h</sup> | 85 (65.9) |

N/A, not applicable; TNM, tumor–node–metastasis.
<sup>a</sup>Reasons why biopsy was not taken are, e.g. in emergency setting (obstructive ileus).
<sup>c</sup>A right-sided tumor is defined as a colon carcinoma in the cecum, colon ascendens, flexura hepatica, or colon transversum.
<sup>d</sup>Different versions of the Union for International Cancer Control (UICC) TNM classification were used. Here, all variables are converted to the UICC TNM version 5 (1997).
<sup>e</sup>Differentiation grade is variously registered as separate or combined subgroups; this is then categorized into combined grades, i.e. well–moderate or poor–undifferentiated.
<sup>f</sup>Chemotherapy regimens included capecitabine monotherapy, capecitabine with oxaliplatin (CAPOX), fluorouracil (5FU) monotherapy and 5FU with leucovorin and oxaliplatin (FOLFOX)
<sup>g</sup>Shown as percentage of total number of recurrence.
<sup>h</sup>Shown as percentage of total number of deaths.

### Self-supervised training of HPL-PanColon and construction of a colorectal histomorphological phenotype atlas

We developed a self-supervised histomorphological phenotype atlas spanning colorectal adenomas and invasive colorectal cancer using the Histomorphological Phenotype Learning framework (28), which integrates self-supervised representation learning with graph-based community detection to identify recurrent tissue phenotypes from unannotated whole-slide images (Fig. 1b). Whole-slide images from the TCGA-COAD, AVANT and ADENOMAS cohorts were tiled at 20× magnification into 224 × 224 pixel patches (0.504 μm per pixel). To assemble a self-supervised training set of comparable scale to the original HPL implementation, 100 tiles were randomly sampled per slide, yielding 83,135 tiles from ADENOMAS, 118,000 from AVANT and 43,403 from TCGA-COAD, for a total of approximately 245,000 tiles. A Barlow Twins encoder (41) was trained on this tile set to learn 128-dimensional embeddings that capture histomorphological variation by maximizing agreement between augmented views of the same tile while minimizing redundancy across embedding features. The resulting embeddings were first overclustered using Leiden community detection (42) at resolution of 5.0 to identify and exclude clusters enriched for technical artifacts. Clustering was then repeated on the filtered embeddings across multiple resolutions, and resolution 2.5 was selected based on the optimal silhouette score. The resulting clusters were defined as histomorphological phenotype clusters (HPCs) and comprised the phenotype atlas used in downstream analyses (Fig. 1b). A 1024-dimensional embedding configuration was also evaluated, but did not improve phenotype clustering quality or downstream performance; therefore, the 128-dimensional representation was retained for subsequent analyses.

### Comparison with existing pathology foundation models

To assess whether a colorectal-neoplasia-specific embedding model was better suited for this study, we compared HPL-PanColon with two existing pathology foundation models, TITAN (32) and UNI (31). In contrast to these broad multi-tissue models, HPL-PanColon was trained on colorectal cohorts explicitly spanning adenomas and invasive colorectal cancer. Tile-level embeddings were generated for tiles from the HPL-PanColon development cohorts and from the UNITED cohort using each model. For the development cohorts, UMAP projections were colored by source cohort, including TCGA-COAD, AVANT, and the internal ADENOMAS cohort, to qualitatively assess cohort-associated structure. For the UNITED cohort, UMAP projections were colored by contributing institution to qualitatively assess institution-associated structure. These analyses were used as visual comparisons of dataset- and institution-associated separation across embedding spaces and supported the use of HPL-PanColon for downstream HPC discovery and cross-institutional survival modelling.

### Survival modeling using SurvCLAM

For downstream survival analysis, we applied HPL-PanColon to the UNITED cohort to generate tile-level embeddings, which were subsequently used as input to a survival-adapted version of CLAM (43), hereafter termed SurvCLAM (Fig. 1c). The objective of this analysis was to predict DFS. The model retained the attention-based multiple instance learning framework of CLAM, in which tile-level features h*_ik_*were aggregated into a patient-level representation

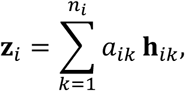

where *a_ik_*denotes the learned attention weight for tile *k*in patient *i*. The aggregated representation was passed through a single-output linear risk head to generate a continuous patient-level DFS risk score, the Colon Histomorphology Prognostic Score (CHiPS). SurvCLAM was trained by minimizing the Cox partial likelihood loss,

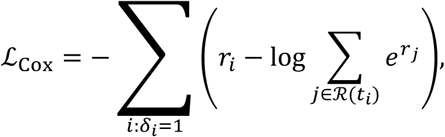

where *r_i_* denotes the predicted CHiPS for patient *i*, *t_i_* the observed follow-up time, *δ_i_* the DFS event indicator, and ℛ(*t_i_*) the risk set at time *t_i_*.

Training and inference were performed at the patient level, such that all slides from an individual patient were grouped into a single bag. The model operated on 128-dimensional input embeddings and used a dropout rate of 0.25.

To evaluate cross-institutional generalizability, SurvCLAM was trained in a leave-one-institution-out design across the UNITED cohort. In each split, one institution was held out for testing, whereas the remaining institutions were used for model development. An internal validation set was created by reserving 20% of patients and institutions from the development cohort. Training was performed with a learning rate of 2 × 10^-4^, weight decay of 1 × 10^-5^, dropout of 0.25, gradient clipping at 1.0 and a maximum of 200 epochs. Learning rate scheduling used reduction on plateau with a factor of 0.5 and patience of 4 epochs.

Performance was summarized from pooled out-of-fold predictions generated on the held-out institution across all splits. Prognostic discrimination for DFS was quantified using the concordance index, and pooled CHiPS values were further stratified into tertiles to generate Kaplan– Meier curves for low-, intermediate- and high-risk groups (Fig. 1d).

### Model interpretability and association of histomorphological phenotypes with risk

To interpret the morphological patterns prioritized by the survival model, we analyzed the SurvCLAM models used during cross-validation. For each held-out patient, CHiPS was derived using the corresponding fold-specific model, ensuring that predictions were generated only for patients unseen during training. Tile-level attention scores were extracted from the final attention layer before patient-level aggregation, thereby providing a measure of the relative contribution of individual tiles to patient-level risk prediction.

To relate model attention to the histomorphological phenotype atlas, we mapped tile embeddings from the UNITED cohort onto the previously defined HPL-PanColon HPCs using k-nearest-neighbor assignment in the learned embedding space (Fig. 1d). This enabled each tile in the UNITED cohort to be assigned to a corresponding HPC, allowing direct comparison between the phenotypes emphasized by the model and the phenotype atlas derived during self-supervised training. We visualized the distribution of UNITED tile embeddings using uniform manifold approximation and projection (UMAP), with tiles colored by assigned HPC to assess the organization of histomorphological phenotypes in the embedding space. We further overlaid tile-level attention scores on the same UMAP representation to identify regions of the phenotype atlas preferentially emphasized by SurvCLAM during risk prediction.

We then quantified the relationship between HPC composition and survival risk using tile-level attention scores and patient-level risk groups derived from out-of-fold CHiPS. For each HPC, we evaluated its representation among highly attended tiles and compared its relative abundance between high- and low-risk patients. In this interpretability analysis, tiles were restricted to the top 25% of attention scores. For each cluster, the fraction of top-attention tiles was computed together with the patient-level distribution of that cluster among high- and low-risk groups. Associations with risk were summarized as the difference in median cluster fraction between high- and low-risk patients, and statistical significance was assessed using a two-sided Mann–Whitney U test followed by Benjamini–Hochberg correction. This analysis was used to identify HPCs preferentially associated with unfavorable or favorable risk states and to determine which phenotypes were most strongly prioritized by SurvCLAM.

Representative tiles from risk-associated HPCs were reviewed by a pathologist, who assigned descriptive histomorphological labels to each cluster on the basis of their predominant architectural and stromal features.

### Association of clinical variables with risk

To examine the relationship between CHiPS and standard clinicopathological factors, we analyzed the distribution of out-of-fold CHiPS values across available clinical and pathological variables, including sex, pT category, pN category and tumor stroma ratio. Continuous variables were compared using two-sided Mann–Whitney U tests or Kruskal–Wallis tests when more than two groups were assessed, whereas categorical variables were compared using chi-squared tests or Fisher’s exact tests, as appropriate. Associations were summarized using differences in group distributions, and p-values were adjusted using the Benjamini–Hochberg procedure when multiple variables were tested. We then evaluated whether CHiPS provided prognostic information independent of established clinicopathological variables by fitting a multivariable Cox proportional hazards model for disease-free survival that included CHiPS, age, sex, pT category, pN category and tumor stroma ratio. To assess the added prognostic value of CHiPS, we compared the discriminative performance of clinical-only, CHiPS-only and combined clinical-plus-CHiPS models using the concordance index. Statistical significance for differences in c-index was assessed by patient-level bootstrap resampling across 1,000 iterations. This analysis was used to determine whether CHiPS captured histomorphological risk information beyond conventional clinicopathological features and improved prognostic stratification when integrated with standard clinical variables.

### Spatial transcriptomic characterization of risk-associated histomorphological phenotypes

In order to define the molecular and cellular correlates of risk-associated histomorphological phenotypes, we applied HPL-PanColon to Visium HD H&E whole-slide images, generated tile-level embeddings and assigned histomorphological phenotype clusters. These phenotype labels were then transferred to 8-μm spatial bins, enabling joint analysis of morphology and spatial transcriptomics.

Cellular context was characterized using the spot-level deconvolution and cell-state annotations provided with the Visium HD dataset. Fine-grained annotations were collapsed into broader biological compartments, including tumor epithelial, fibroblast/stromal, smooth muscle/perivascular, endothelial, myeloid, B/plasma, T/NK, intestinal epithelial, neuronal, and other categories. For each HPC, enrichment of each compartment was quantified relative to its overall frequency across all assigned bins and summarized as log2 fold-enrichment values. To further characterize the molecular programs associated with each phenotype, differential gene expression analysis was performed for each HPC using the assigned spatial bins, and the resulting HPC-associated genes were analyzed by over-representation analysis (ORA) using the MSigDB Hallmark 2020 pathway collection (44).

## Data availability

The publicly available TCGA-COAD can be accessed at Genomic Data Commons portal (https://gdc.cancer.gov/). The AVANT data that support the findings of this study are available from Genentech Inc., Roche. However, access to these data is restricted as they were used under license for the current study and are not for commercial use to protect patient privacy. Data may however be available from the authors upon request for research purposes with permission from Genentech Inc., Roche. The ADENOMAS cohort from NYU and the UNITED cohort are not publicly available because of institutional privacy restrictions. Python software packages and codes are available from our previous publications (27, 28). The remaining data are available within the article and supplementary information.

## Code availability

Code used for survival modeling and downstream analysis is available at https://github.com/hortensele/SurvCLAM and https://github.com/hortensele/PanColon-CHiPS.

## Acknowledgement

We thank the team of NYU Langone High Performance Computing (HPC) Core’s resources supporting us to perform the analysis. We would like to thank the NYU Applied Bioinformatics Laboratories (ABL) for providing bioinformatics support and helping with analysis of the data.

## Author contributions

H.L. executed data processing, model training and data analysis. F.H. and W.E.M. provided H&E slides and corresponding clinical data from the UNITED study. S.H. and D.C. provided H&E slides for the ADENOMAS cohort. F.H. and A.K. provided histological assessment and interpretation of the tiles. F.H. and H.L. wrote the first draft of the manuscript which was later revised and approved by all co-authors. H.L., N.C. and A.T. provided expertise in deep learning, biostatistics and bioinformatics. F.H., A.K., K.C.M.J.P., S.H. and W.E.M. provided expertise in colon cancer histopathology and biology. A.T. supervised the study.

## References

1. Siegel RL, Miller KD, Wagle NS, Jemal A. Cancer statistics, 2023. CA Cancer J Clin. 2023;73(1):17–48.

2. Brierley J, Gospodarowicz MK, Wittekind C. TNM classification of malignant tumours. Eighth edition. ed. Chichester, West Sussex, UK ;: John Wiley C Sons, Inc.; 2017.

3. Argilés G, Tabernero J, Labianca R, Hochhauser D, Salazar R, Iveson T, et al. Localised colon cancer: ESMO Clinical Practice Guidelines for diagnosis, treatment and follow-up. Ann Oncol. 2020;31(10):1291–305.

4. Baxter NN, Kennedy EB, Bergsland E, Berlin J, George TJ, Gill S, et al. Adjuvant Therapy for Stage II Colon Cancer: ASCO Guideline Update. J Clin Oncol. 2022;40(8):892–910.

5. Weiser MR. AJCC 8th Edition: Colorectal Cancer. Ann Surg Oncol. 2018;25(6):1454–5.

6. Brierley JD, Giuliani M, O’Sullivan B, Rous B, Eycken EV. TNM Classification of Malignant Tumours, 9th Edition 2025. 272 p.

7. Federatie Medisch Specialisten. Richtlijn Colorectaal Carcinoom 2020 [cited 2025 13-01-2025]. Available from: https://richtlijnendatabase.nl/richtlijn/colorectaal_carcinoom_crc/startpagina_-_crc.html

8. Koncina E, Haan S, Rauh S, Letellier E. Prognostic and Predictive Molecular Biomarkers for Colorectal Cancer: Updates and Challenges. Cancers (Basel). 2020;12(2).

9. Auclin E, Zaanan A, Vernerey D, Douard R, Gallois C, Laurent-Puig P, et al. Subgroups and prognostication in stage III colon cancer: future perspectives for adjuvant therapy. Annals of Oncology. 2017;28(5):958–68.

10. Sobrero AF, Puccini A, Shi Q, Grothey A, Andrè T, Shields AF, et al. A new prognostic and predictive tool for shared decision making in stage III colon cancer. Eur J Cancer. 2020;138:182–8.

11. Gomez D, Calderón C, Carmona-Bayonas A, Cacho Lavin D, Muñoz MM, Martinez Cabañez R, et al. Impact of adjuvant therapy toxicity on quality of life and emotional symptoms in patients with colon cancer: a latent class analysis. Clin Transl Oncol. 2021;23(3):657–62.

12. Lugli A, Kirsch R, Ajioka Y, Bosman F, Cathomas G, Dawson H, et al. Recommendations for reporting tumor budding in colorectal cancer based on the International Tumor Budding Consensus Conference (ITBCC) 2016. Mod Pathol. 2017;30(9):1299–311.

13. Roth AD, Delorenzi M, Tejpar S, Yan P, Klingbiel D, Fiocca R, et al. Integrated analysis of molecular and clinical prognostic factors in stage II/III colon cancer. J Natl Cancer Inst. 2012;104(21):1635–46.

14. Fatemi N, Mirbahari SN, Tierling S, Sanjabi F, Shahrivari S, AmeliMojarad M, et al. Emerging Frontiers in Colorectal Cancer Therapy: From Targeted Molecules to Immunomodulatory Breakthroughs and Cell-Based Approaches. Dig Dis Sci. 2025;70(3):919–42.

15. Anderson NM, Simon MC. The tumor microenvironment. Curr Biol. 2020;30(16):R921–r5.

16. van Pelt GW, Kjær-Frifeldt S, van Krieken J, Al Dieri R, Morreau H, Tollenaar R, et al. Scoring the tumor-stroma ratio in colon cancer: procedure and recommendations. Virchows Arch. 2018;473(4):405–12.

17. Mesker WE, Junggeburt JM, Szuhai K, de Heer P, Morreau H, Tanke HJ, et al. The carcinoma-stromal ratio of colon carcinoma is an independent factor for survival compared to lymph node status and tumor stage. Cell Oncol. 2007;29(5):387–98.

18. Mesker WE, Liefers GJ, Junggeburt JM, van Pelt GW, Alberici P, Kuppen PJ, et al. Presence of a high amount of stroma and downregulation of SMAD4 predict for worse survival for stage I-II colon cancer patients. Cell Oncol. 2009;31(3):169–78.

19. Bera K, Schalper KA, Rimm DL, Velcheti V, Madabhushi A. Artificial intelligence in digital pathology - new tools for diagnosis and precision oncology. Nat Rev Clin Oncol. 2019;16(11):703–15.

20. Tolkach Y, Dohmgörgen T, Toma M, Kristiansen G. High-accuracy prostate cancer pathology using deep learning. Nature Machine Intelligence. 2020;2(7):411–8.

21. Campanella G, Hanna MG, Geneslaw L, Miraflor A, Werneck Krauss Silva V, Busam KJ, et al. Clinical-grade computational pathology using weakly supervised deep learning on whole slide images. Nature Medicine. 2019;25(8):1301–9.

22. Coudray N, Ocampo PS, Sakellaropoulos T, Narula N, Snuderl M, Fenyö D, et al. Classification and mutation prediction from non-small cell lung cancer histopathology images using deep learning. Nat Med. 2018;24(10):1559–67.

23. Heilijgers F, Polack M, Roodvoets AGH, Meershoek - Klein Kranenbarg E, Peeters KCMJ, Buettner R, et al. Automating tumor–stroma ratio quantification in colon cancer patients from the UNITED study. ESMO Open. 2026;11(1):105934.

24. Bilal M, Raza SEA, Azam A, Graham S, Ilyas M, Cree IA, et al. Development and validation of a weakly supervised deep learning framework to predict the status of molecular pathways and key mutations in colorectal cancer from routine histology images: a retrospective study. The Lancet Digital Health. 2021;3(12):e763–e72.

25. Wulczyn E, Steiner DF, Moran M, Plass M, Reihs R, Tan F, et al. Interpretable survival prediction for colorectal cancer using deep learning. npj Digital Medicine. 2021;4(1):71.

26. Rani V, Nabi ST, Kumar M, Mittal A, Kumar K. Self-supervised Learning: A Succinct Review. Arch Comput Methods Eng. 2023;30(4):2761–75.

27. Liu B, Polack M, Coudray N, Claudio Quiros A, Sakellaropoulos T, Le H, et al. Self-supervised learning reveals clinically relevant histomorphological patterns for therapeutic strategies in colon cancer. Nature Communications. 2025;16(1):2328.

28. Claudio Quiros A, Coudray N, Yeaton A, Yang X, Liu B, Le H, et al. Mapping the landscape of histomorphological cancer phenotypes using self-supervised learning on unannotated pathology slides. Nature Communications. 2024;15(1):4596.

29. Chen C, Lu MY, Williamson DFK, Chen TY, Schaumberg AJ, Mahmood F. Fast and scalable search of whole-slide images via self-supervised deep learning. Nat Biomed Eng. 2022;6(12):1420–34.

30. Xu H, Usuyama N, Bagga J, Zhang S, Rao R, Naumann T, et al. A whole-slide foundation model for digital pathology from real-world data. Nature. 2024;630(8015):181–8.

31. Chen RJ, Ding T, Lu MY, Williamson DFK, Jaume G, Song AH, et al. Towards a general-purpose foundation model for computational pathology. Nat Med. 2024;30(3):850–62.

32. Ding T, Wagner SJ, Song AH, Chen RJ, Lu MY, Zhang A, et al. A multimodal whole-slide foundation model for pathology. Nat Med. 2025;31(11):3749–61.

33. Polack M, Smit MA, van Pelt GW, Roodvoets AGH, Meershoek-Klein Kranenbarg E, Putter H, et al. Results from the UNITED study: a multicenter study validating the prognostic effect of the tumor–stroma ratio in colon cancer. ESMO Open. 2024;9(4):102988.

34. van Pelt GW, Sandberg TP, Morreau H, Gelderblom H, van Krieken J, Tollenaar R, et al. The tumour-stroma ratio in colon cancer: the biological role and its prognostic impact. Histopathology. 2018;73(2):197–206.

35. Zunder S, van der Wilk P, Gelderblom H, Dekker T, Mancao C, Kiialainen A, et al. Stromal organization as predictive biomarker for the treatment of colon cancer with adjuvant bevacizumab; a post-hoc analysis of the AVANT trial. Cell Oncol (Dordr). 2019;42(5):717–25.

36. Strous MTA, Faes TKE, Gubbels ALHM, van der Linden RLA, Mesker WE, Bosscha K, et al. A high tumour-stroma ratio (TSR) in colon tumours and its metastatic lymph nodes predicts poor cancer-free survival and chemo resistance. Clinical and Translational Oncology. 2022;24(6):1047–58.

37. Zunder SM, van Pelt GW, Gelderblom HJ, Mancao C, Putter H, Tollenaar RA, et al. Predictive potential of tumour-stroma ratio on benefit from adjuvant bevacizumab in high-risk stage II and stage III colon cancer. British Journal of Cancer. 2018;119(2):164–9.

38. de Gramont A, Van Cutsem E, Schmoll H-J, Tabernero J, Clarke S, Moore MJ, et al. Bevacizumab plus oxaliplatin-based chemotherapy as adjuvant treatment for colon cancer (AVANT): a phase 3 randomised controlled trial. The Lancet Oncology. 2012;13(12):1225–33.

39. TCGA Colon Adenocarcinoma (TCGA-COAD). [Internet]. Available from: https://portal.gdc.cancer.gov/projects/TCGA-COAD.

40. Oliveira MFd, Romero JP, Chung M, Williams SR, Gottscho AD, Gupta A, et al. High-definition spatial transcriptomic profiling of immune cell populations in colorectal cancer. Nature Genetics. 2025;57(6):1512–23.

41. Zbontar J, Jing L, Misra I, LeCun Y, Deny S. Barlow Twins: Self-Supervised Learning via Redundancy Reduction. CoRR. 2021;abs/2103.03230.

42. Traag VA, Waltman L, van Eck NJ. From Louvain to Leiden: guaranteeing well-connected communities. CoRR. 2018;abs/1810.08473.

43. Lu MY, Williamson DFK, Chen TY, Chen RJ, Barbieri M, Mahmood F. Data-efficient and weakly supervised computational pathology on whole-slide images. Nature Biomedical Engineering. 2021;5(6):555–70.

44. Liberzon A, Birger C, Thorvaldsdóttir H, Ghandi M, Mesirov JP, Tamayo P. The Molecular Signatures Database (MSigDB) hallmark gene set collection. Cell Syst. 2015;1(6):417–25.

