## Supplemental Figures for "Multicenter self-supervised computational pathology identifies prognostic histomorphological phenotypes in colorectal cancer"

**[#]UNITED collaboration**

F. Heilijgers, M. Polack, A. G. H. Roodvoets, E. Meershoek-Klein Kranenbarg, K. C. M. J. Peeters, W. E. Mesker, G. Petrushevska, M. Bogdanovska Todorovska, P. Zdravkoski, P. Karagjozov, D. Dzamba, S. Antovic, S. Kostadinova Kunovska, E. M. V. de Cuba, F. Beverdam, J. Jansen, M. Vermaas, G. Gašljević, S. Kjær-Frifeldt, J. Lindeberg, M. Strous, N. W. J. Bulkmans, L. Mekenkamp, R. Hoekstra, M. Sie, M. Cuatrecasas, S. Simonetti, M. Teresa Rodrigo, I. Archilla Sanz, N. E Leeuwis-Fedorovich, K. A Talsma, R. M. Souza da Silva, M. M. Lacle, M. Koopman, J. W. T. Dekker, A. van Tilburg, P. Nuciforo, X. Villalobos Alberú, S. Landolfi, A. Zucchiatti, E. Witteveen, A. Bordbar, M. P. Hendriks, L. Arensman, S. Natu, R. A. E. M. Tollenaar, M. A. Smit, G. W. van Pelt, H. Putter, A. S. L. P. Crobach, H. Gelderblom, M. Fontes E Sousa, P. Borralho Nunes, J. Cruz, A. Raimundo, M. J. Brito, V. Terpstra, L. M. Zakhartseva, R. Al Dieri, R. Feakins, E. Dequeker, J. Roodhart, J. H. J. M. van Krieken, K. Kerckhoffs

[1]Department of Surgery, Leiden University Medical Center, Leiden, The Netherlands

[2]Division of Precision Medicine, Department of Medicine, Grossman School of Medicine, New York University, New York, New York

[3]Department of Pathology, New York University Grossman School of Medicine, New York, NY, USA

[4]Applied Bioinformatics Laboratories, New York University Grossman School of Medicine, New York, NY, USA

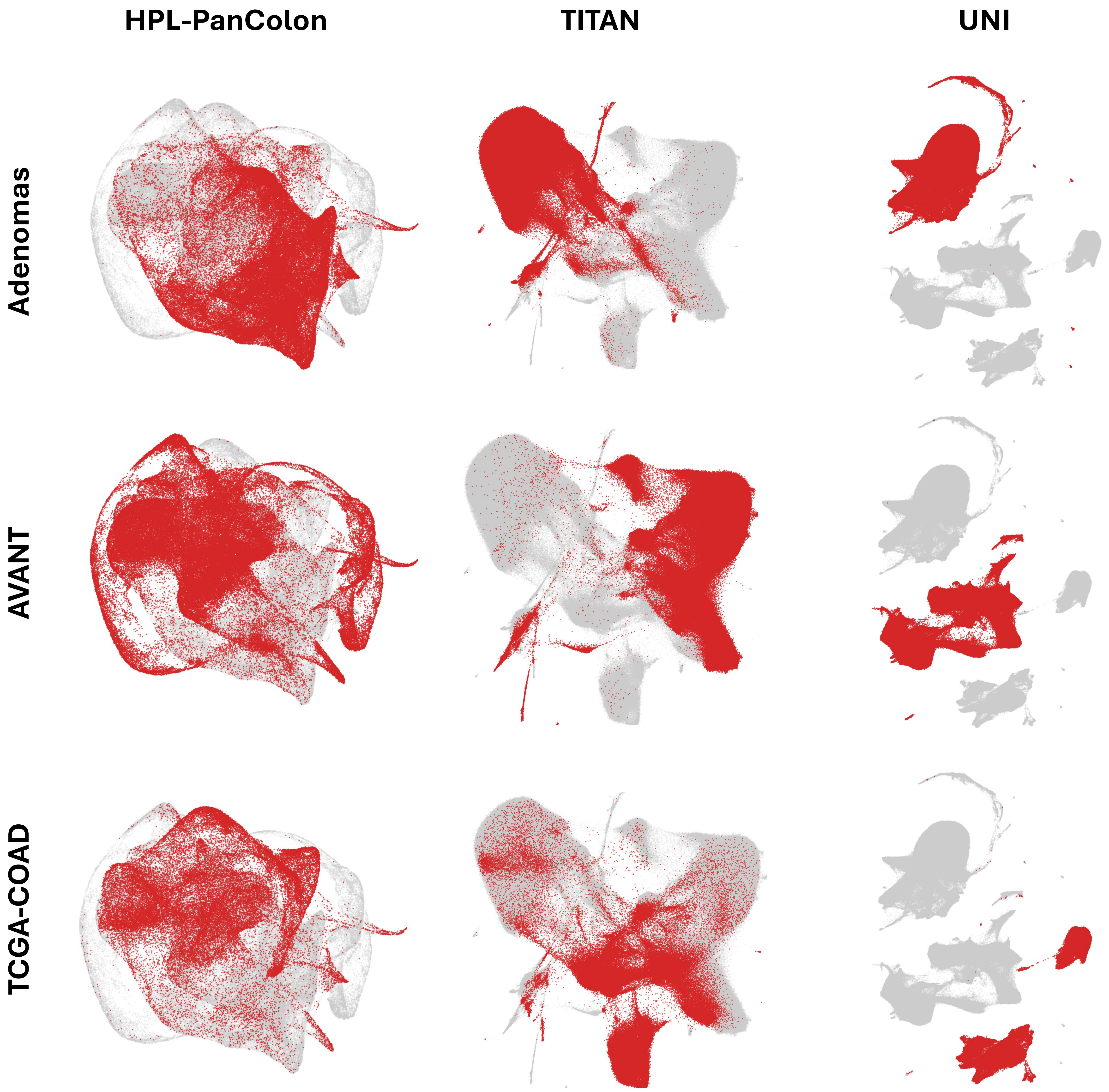


**Supplementary Figure 1 | Comparison of cohort-associated structure across pathology embedding models in the HPL-PanColon development set.**
UMAP projections of tile embeddings generated by HPL-PanColon, TITAN, and UNI across the HPL-PanColon development cohorts, including TCGA-COAD, AVANT, and ADENOMAS. Points are colored by source cohort. Compared with TITAN and UNI, HPL-PanColon showed greater mixing across development cohorts, supporting the use of a colorectal-neoplasia-specific representation for downstream histomorphological phenotype discovery.

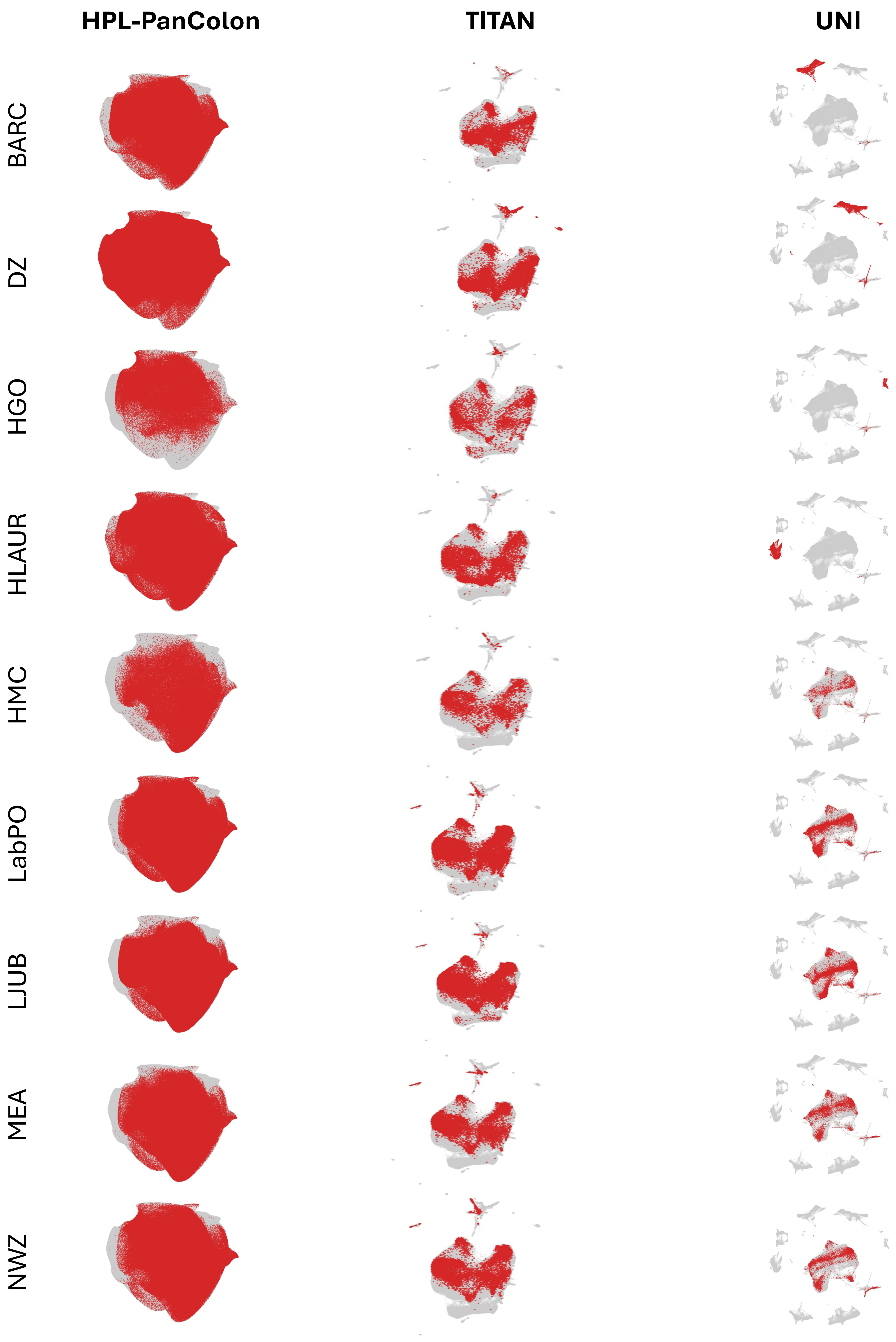


**Supplementary Figure 2 | Comparison of institution-associated structure across pathology embedding models in the UNITED cohort.**
UMAP projections of tile embeddings generated by HPL-PanColon, TITAN, and UNI in the UNITED cohort. Points are colored by contributing institution. HPL-PanColon showed reduced institution-associated separation compared with TITAN and UNI, supporting its use for cross-institutional survival modeling in UNITED.

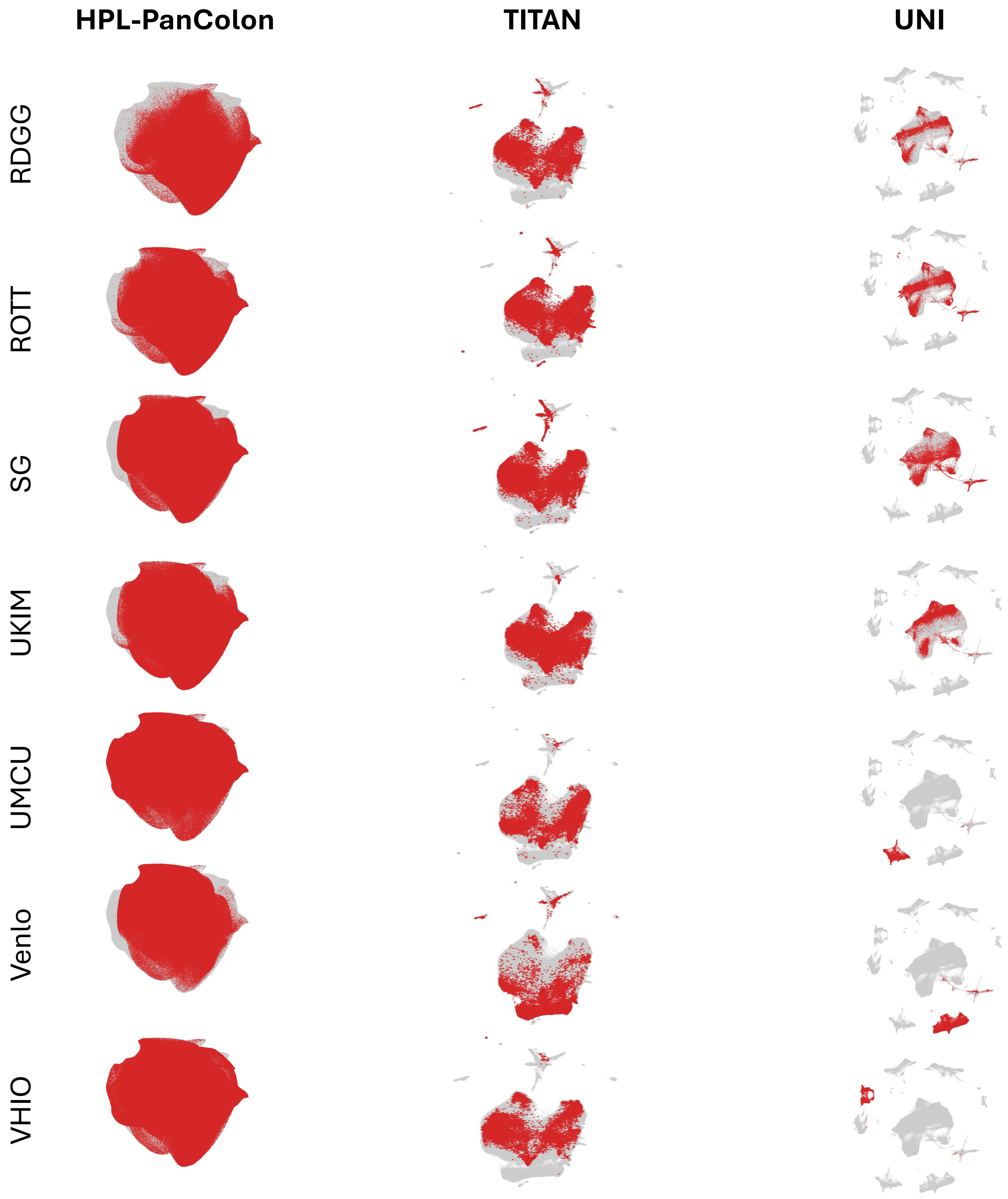


**Supplementary Figure 2 continued.**

HPC0

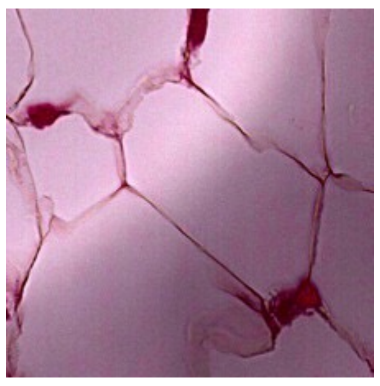

Adipose tissue - pericolorectal/mesorectal adipose tissue with varying proportions of fat lobules, fibrovascular septa, stromal cells

HPC1

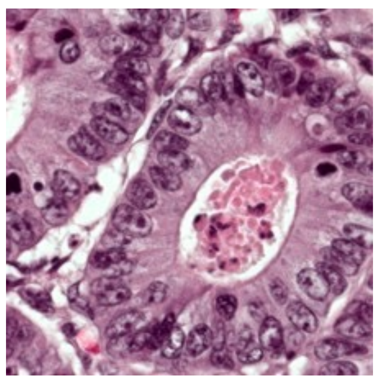

Colorectal Adenocarcinoma - characterized by irregular, crowded glands with hyperchromatic nuclei, desmoplastic stroma, variable architecture, and loss of normal crypt organization.

HPC2

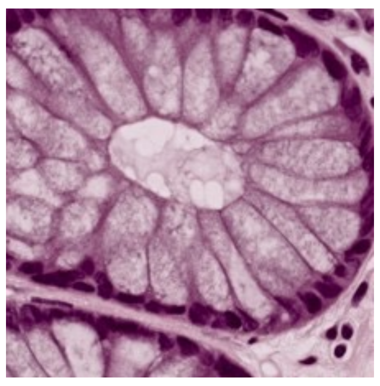

Normal colonic mucosa with well-organized, evenly spaced crypts lined by abundant goblet cells and maintained epithelial polarity on a regular lamina propria background

HPC3

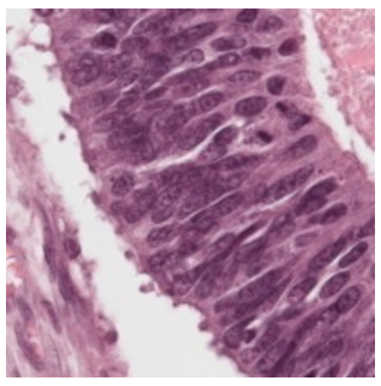

Tubular adenoma with dysplastic epithelium showing elongated, hyperchromatic, pseudostratified nuclei, loss of goblet cells, and crowded tubular glands with preserved but architecturally distorted crypt structure.

HPC4

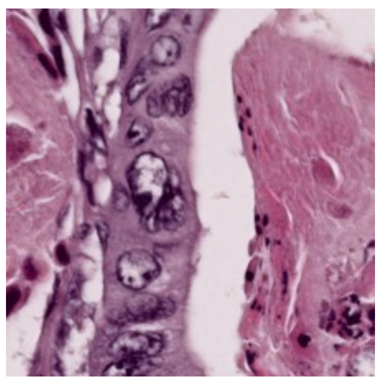

Colorectal adenocarcinoma- Infiltrating malignant glands embedded in prominent desmoplastic stroma with thick-walled vessels

HPC5

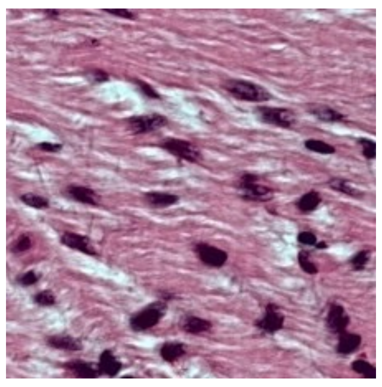

Normal colonic muscularis propria showing interlacing bundles of smooth muscle cells with elongated bland nuclei.

HPC6

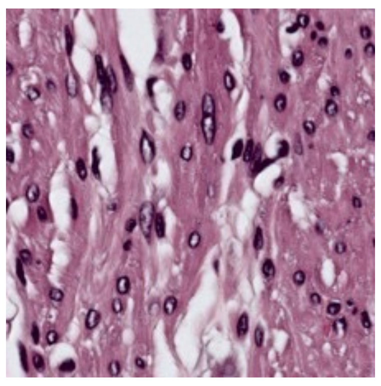

Colonic muscularis propria with smooth muscle bundles showing slightly increased nuclear variability, scattered inflammatory cells, and focal red cell extravasation, likely representing muscularis propria with mild reactive changes.

HPC7

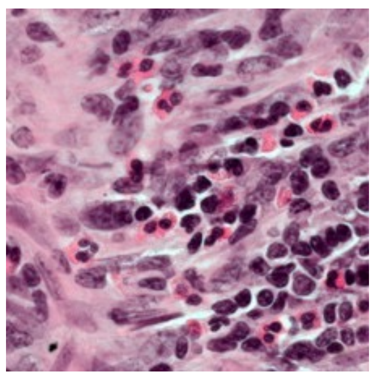

Inflammatory infiltrate composed predominantly lymphocytes and plasma cells with scattered eosinophils and extravasated Rbc's within the colonic wall.

**Supplementary Figure 3 | Pathologist-annotated reference atlas of HPL-PanColon histomorphological phenotype clusters.**

HPC8

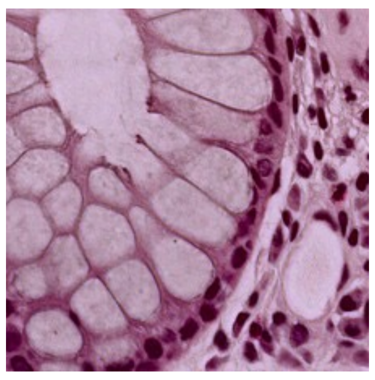

Normal colonic mucosa with well-organized, evenly spaced crypts lined by abundant goblet cells and maintained epithelial polarity on a regular lamina propria background

HPC9

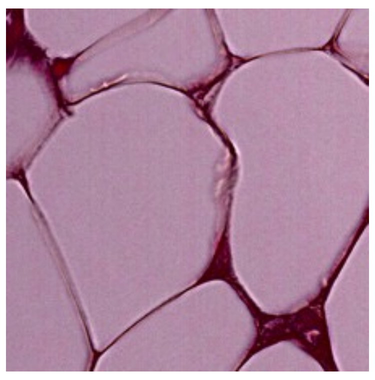

Adipose tissue - pericolorectal/mesorectal adipose tissue with varying proportions of fat lobules, fibrovascular septa, stromal cells

HPC10

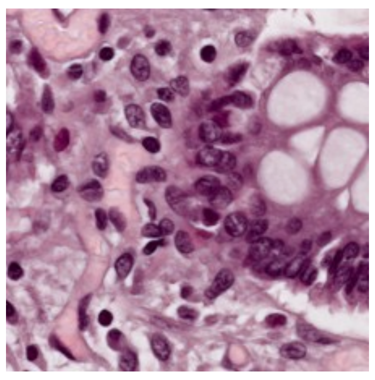

Inflammatory  infiltrate composed predominantly lymphocytes and plasma cells with scattered eosinophils and extravasated Rbc s within the colonic wall.

HPC11

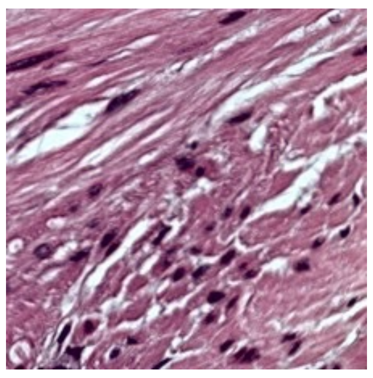

Normal colonic muscularis propria showing interlacing bundles of smooth muscle cells with elongated bland nuclei .

HPC12

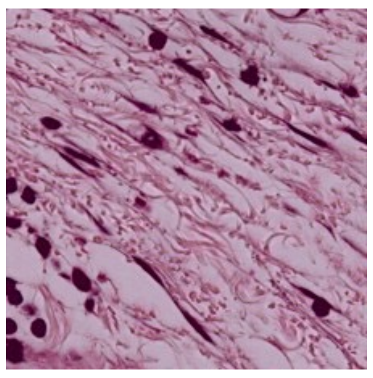

Paucicellular collagenous/fibrous stroma with scattered bland spindle cells, small vessels, and rare inflammatory cells, consistent with desmoplastic stroma

HPC13

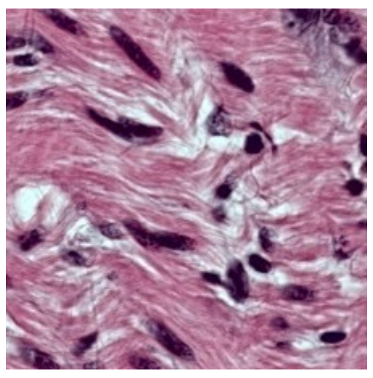

Normal colonic muscularis propria with bland smooth muscle bundles in interlacing orientations, scattered elongated nuclei, and small intervening capillaries.

HPC14

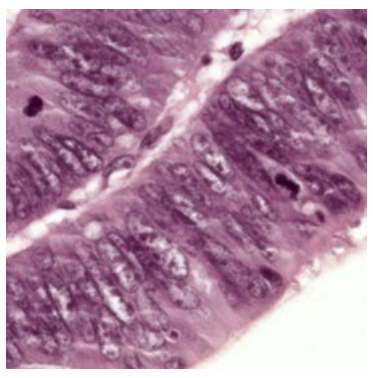

Tubular adenoma fragments with dysplastic columnar epithelium, elongated hyperchromatic pseudostratified nuclei, and dilated irregular glandular lumina set in a pale stromal background

HPC15

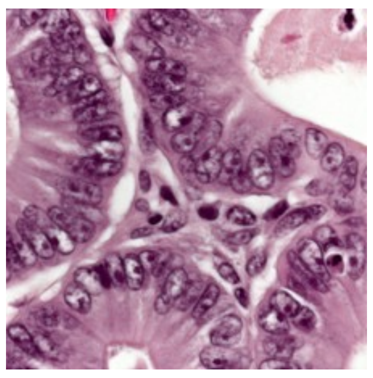

Invasive colorectal adenocarcinoma with irregular malignant glands, hyperchromatic pleomorphic nuclei, prominent nucleoli, and Glandular lumen show necrosis.

**Supplementary Figure 3 continued.**

HPC16

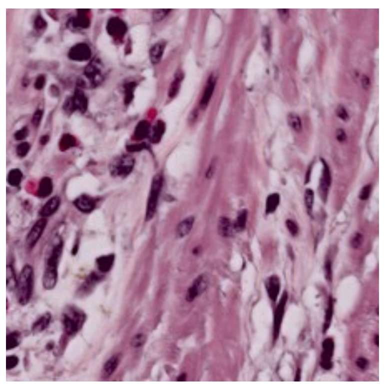

Mixed cluster showing smooth muscle bundles of muscularis propria intimately admixed with sheets of inflammatory cells and scattered hyperchromatic nuclei, suggesting tumor-infiltrating inflammatory response at the invasive front of colorectal carcinoma.

HPC17

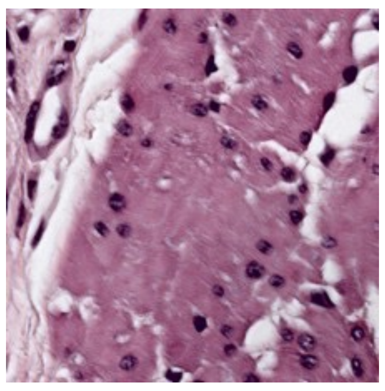

Colonic muscularis propria with interlacing smooth muscle bundles, prominent fibrous septa, thick-walled vessels, and focal nerve fascicles, representing a deeper layer of the colonic wall

HPC18

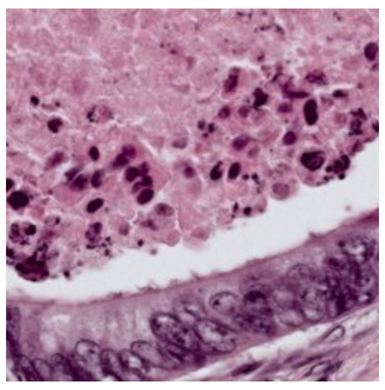

Extensive necrotic debris with ghost cell outlines, karyorrhectic nuclear fragments, and scattered viable tumor cell clusters at the periphery, consistent with tumor necrosis in colorectal carcinoma.

HPC19

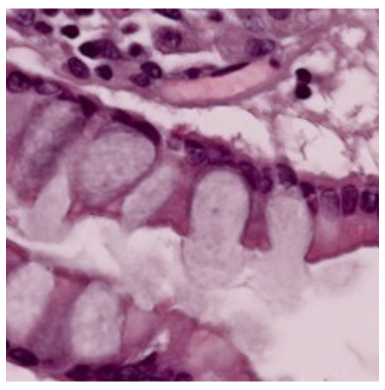

Normal colonic wall

HPC20

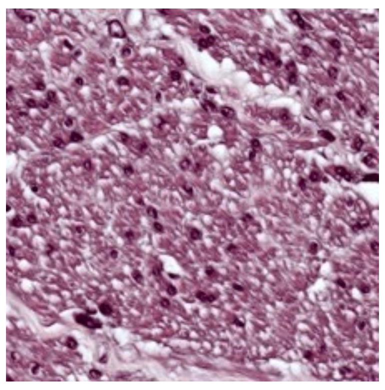

Normal colonic muscularis propria with uniform bland smooth muscle bundles in interlacing orientations, elongated cigar-shaped nuclei, and minimal intervening stroma.

HPC21

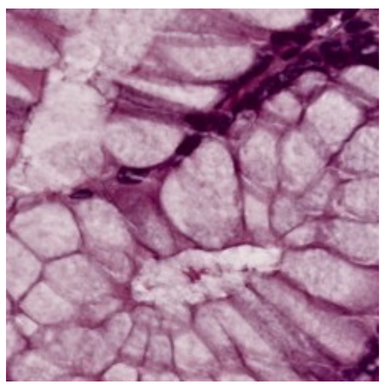

Normal colon

HPC22

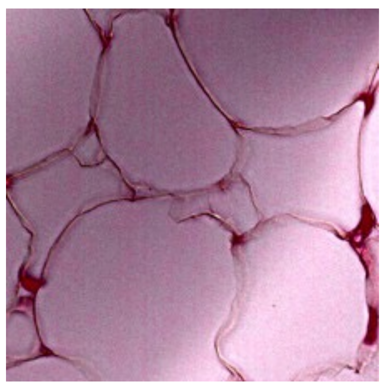

Adipose tissue - pericolorectal/mesorectal adipose tissue with varying proportions of fat lobules

HPC23

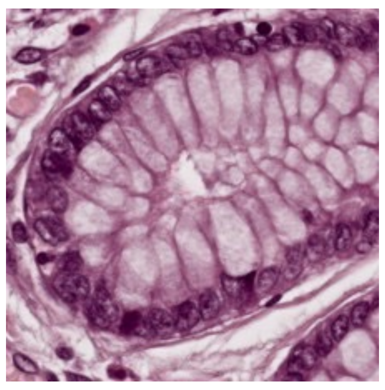

Normal colonic mucosa with well-formed round to oval crypts lined by abundant goblet cells, maintained epithelial polarity, and orderly lamina propria, consistent with normal colon.

**Supplementary Figure 3 continued.**

HPC24

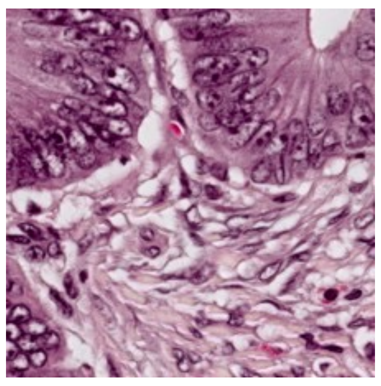

Invasive adenocarcinoma with irregular malignant glands embedded in desmoplastic fibromuscular stroma, showing hyperchromatic nuclei, loss of polarity, and infiltrative growth pattern.

HPC25

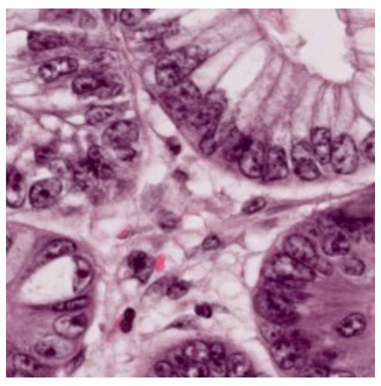

Mixed cluster of normal colonic crypts with goblet cells and dysplastic glands showing hyperchromatic pseudostratified nuclei and reduced goblet cells, consistent with normal mucosa admixed with tubular adenoma fragments.

HPC26

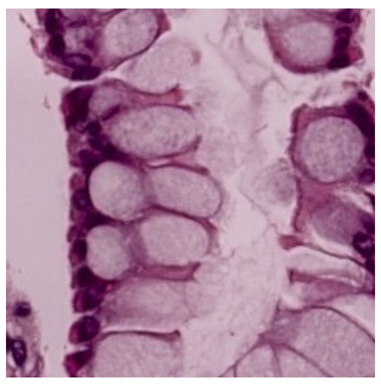

Normal colon

HPC27

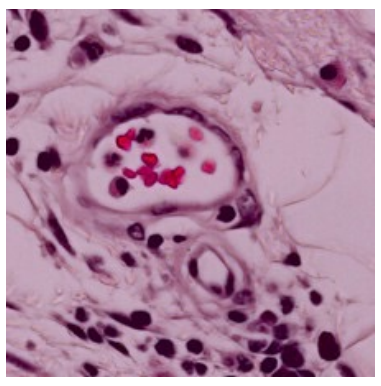

Fibrovascular stroma with prominent thick-walled and thin-walled vessels containing red blood cells, scattered inflammatory cells, and bland fibroblasts, consistent with submucosal or peritumoral vascular-rich connective tissue.

HPC28

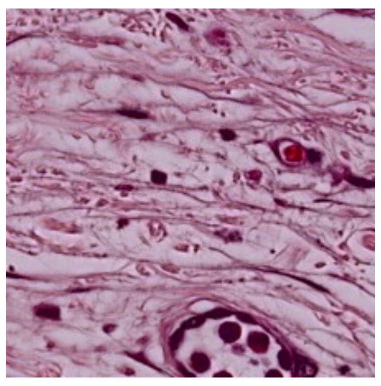

Paucicellular dense collagenous stroma with widely spaced bland fibroblasts, scattered small vessels, and rare inflammatory cells, consistent with pericolorectal or submucosal fibrosis/desmoplastic stroma

HPC29

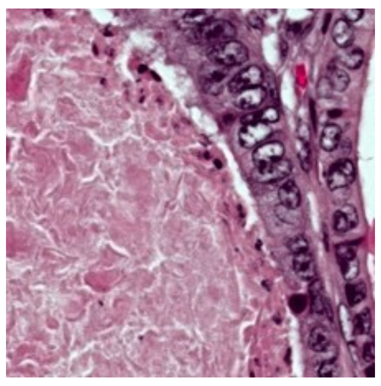

Necrotic debris, and scattered tumor fragments, consistent with invasive tumor front with associated inflammation, necrosis.

HPC30

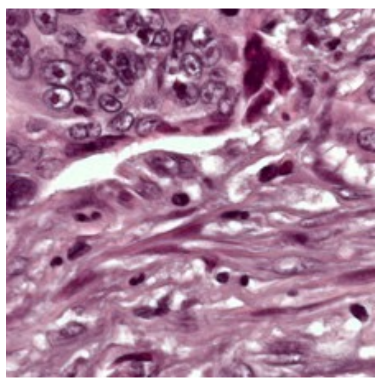

Cellular spindle cell proliferation with interlacing fascicles, mild nuclear pleomorphism, scattered inflammatory cells, and focal red cell extravasation, consistent with desmoplastic stroma or muscularis propria with reactive stromal changes at the tumor interface.

HPC31

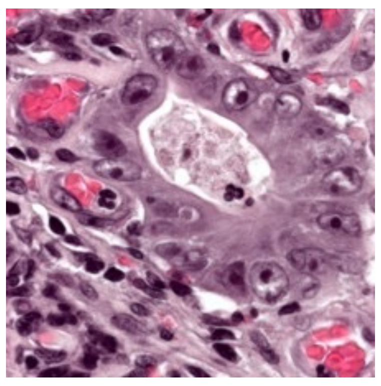

Invasive adenocarcinoma with irregular malignant glands, hyperchromatic pleomorphic nuclei, prominent red cell extravasation, and desmoplastic stroma, consistent with invasive colorectal carcinoma.

**Supplementary Figure 3 continued.**

HPC32

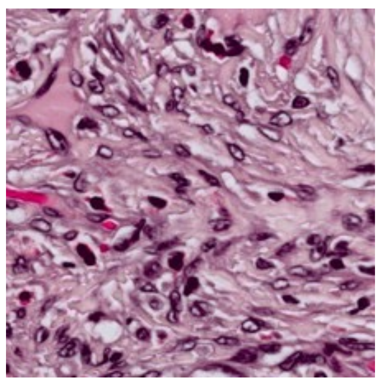

Cellular spindle cell proliferation in interlacing fascicles with mild nuclear pleomorphism, scattered inflammatory cells, and focal vascular spaces, consistent with muscularis propria with reactive stromal changes or desmoplastic tumor stroma.

HPC33

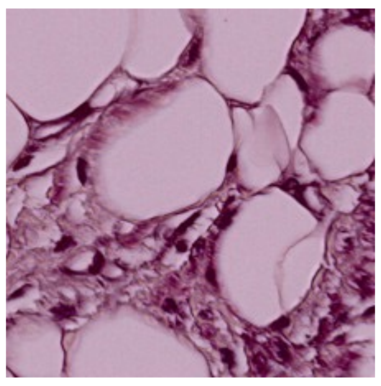

Adipose tissue with large clear adipocyte vacuoles, thin fibrovascular septa, scattered bland fibroblasts, and small vessels, consistent with normal mesorectal fat.

HPC34

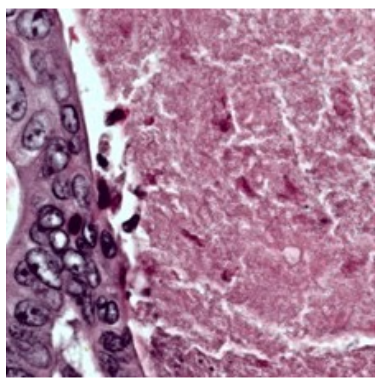

Mainly necrosis with occasioval tile showing tumor

HPC35

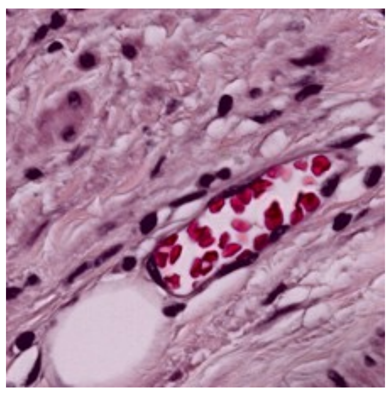

Fibrovascular connective tissue with bland spindle cells, thick collagen bundles, scattered small vessels with red cell extravasation, and focal adipocytes, consistent with pericolorectal or submucosal fibrovascular stroma.

HPC36

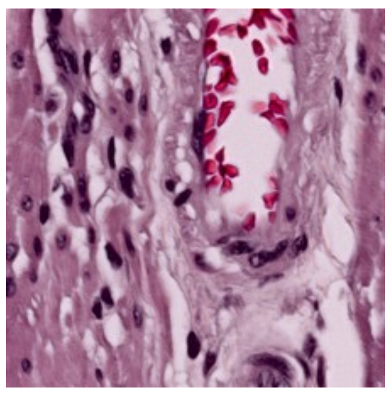

Fibrovascular connective tissue with bland spindle cells, thick collagen bundles, scattered small vessels with red cell extravasation, and focal adipocytes, consistent with pericolorectal or submucosal fibrovascular stroma.

HPC37

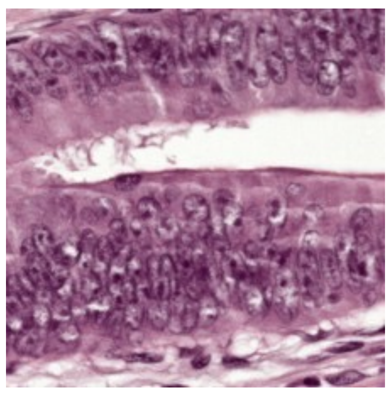

Crowded dysplastic glands with hyperchromatic pseudostratified nuclei, loss of goblet cells, and minimal intervening stroma, consistent with tubular adenoma.

HPC38

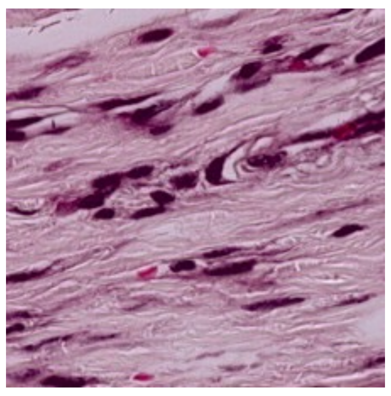

Normal colonic muscularis propria with bland smooth muscle bundles in interlacing orientations, scattered elongated nuclei, and small intervening capillaries.

HPC39

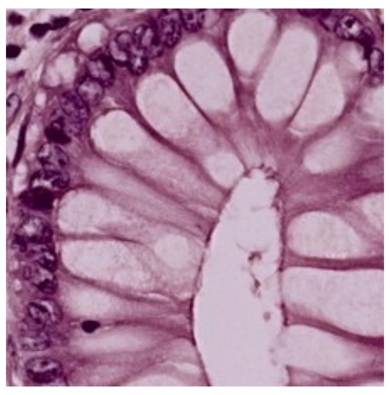

Normal colon.

**Supplementary Figure 3 continued.**

HPC40

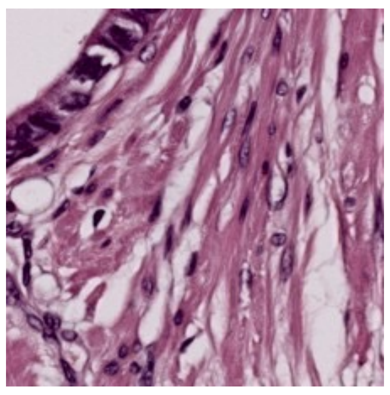

Hypocellular desmoplastic stroma with dense fibrotic collagen bundles, widely spaced bland spindle cells, and minimal visible tumor epithelium, consistent with treatment-related or spontaneous stromal fibrosis surrounding colorectal carcinoma.

HPC41

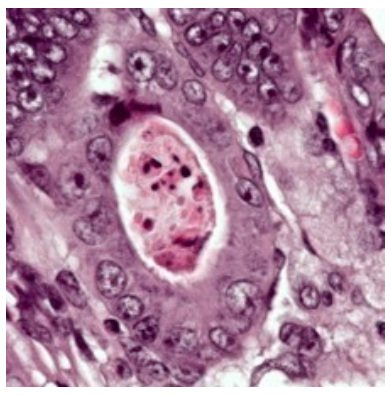

Colorectal adenocarcinoma

HPC42

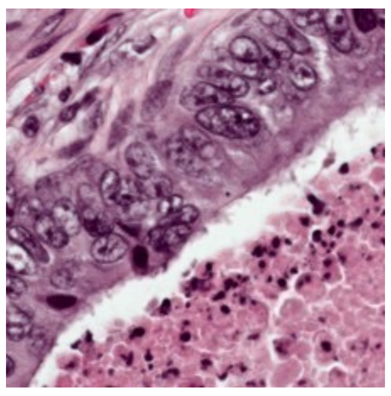

Invasive adenocarcinoma with irregular malignant glands, hyperchromatic pleomorphic nuclei, dilated lumina with intraluminal necrosis, and desmoplastic stromal background, consistent with moderately differentiated colorectal adenocarcinoma.

HPC43

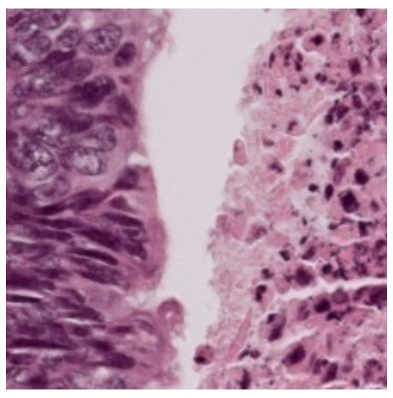

Invasive adenocarcinoma with irregular malignant glands, hyperchromatic pleomorphic nuclei, prominent intraluminal and stromal necrotic debris, and focal red cell extravasation, consistent with moderately to poorly differentiated colorectal carcinoma with necrosis.

HPC44

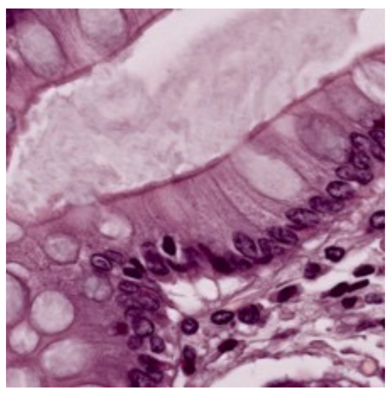

Normal colon with mild hyperplastic change.

HPC45

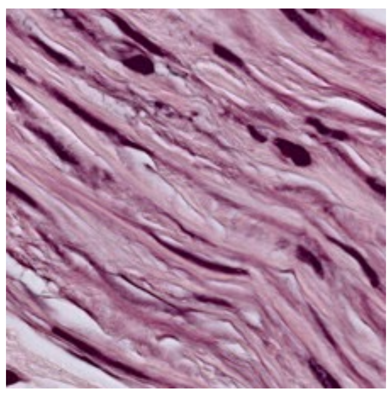

Normal colonic muscularis propria with uniform bland smooth muscle bundles in interlacing orientations, elongated cigar-shaped nuclei, and minimal intervening stroma.

HPC46

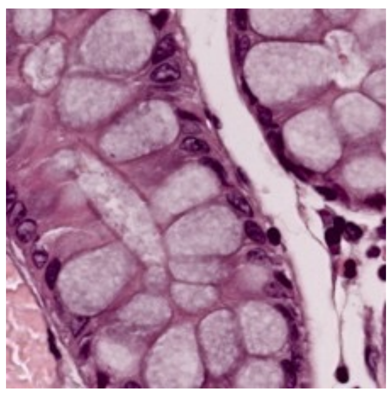

Normal colon

HPC47

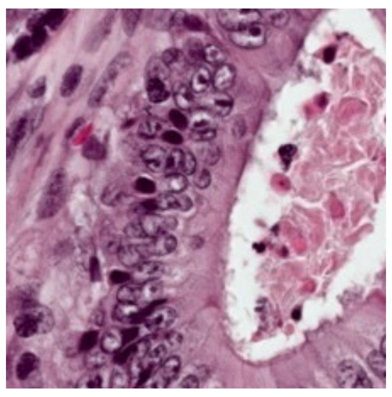

Colonic adenocarcinoma

**Supplementary Figure 3 continued.**

HPC48

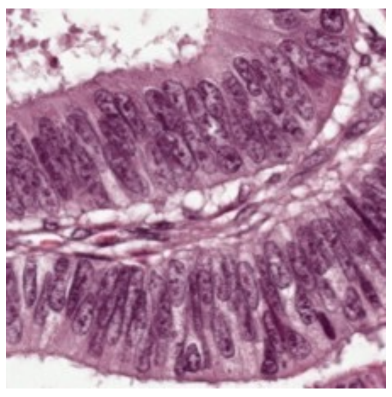

Mainly tubular adenoma fragments with few tiles showing normal colon.

HPC49

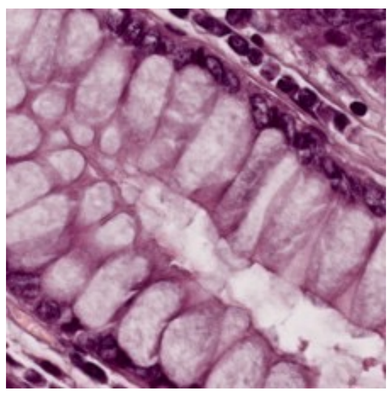

Normal colon.

HPC50

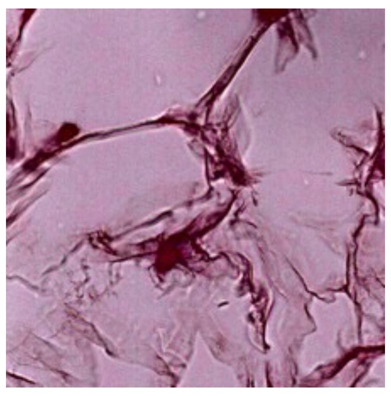

Processing artifacts.

HPC51

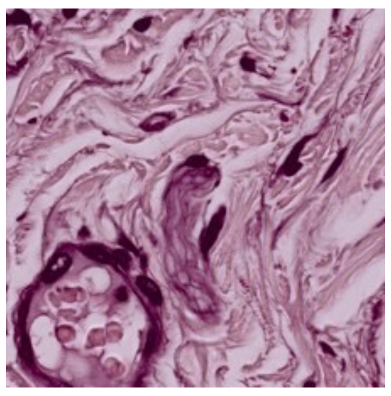

Dense hyalinized collagenous stroma with prominent nerve fascicles, thick-walled vessels, and scattered bland fibroblasts, consistent with perineural fibrosis or submucosal/pericolorectal connective tissue

HPC52

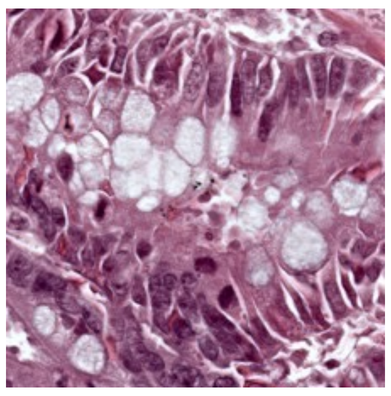

Mixed cluster of normal goblet cell-rich colonic crypts admixed with dysplastic glands showing hyperchromatic pseudostratified nuclei and reduced mucin, consistent with normal mucosa transitioning to tubular adenoma.

HPC53

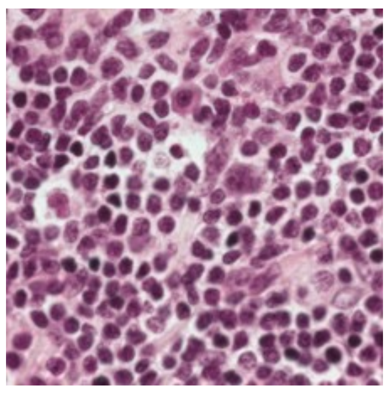

Inflammatory cells with extravasated RBC s.

HPC54

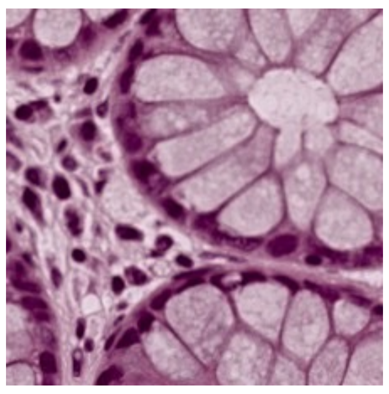

Normal colon

HPC55

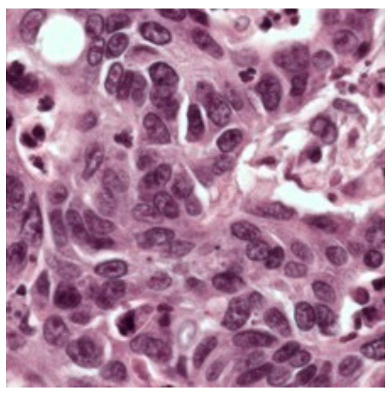

Sheets of poorly differentiated malignant cells with marked nuclear pleomorphism, prominent nucleoli, frequent mitoses, and minimal gland formation, consistent with poorly differentiated colorectal adenocarcinoma

**Supplementary Figure 3 continued.**

HPC56

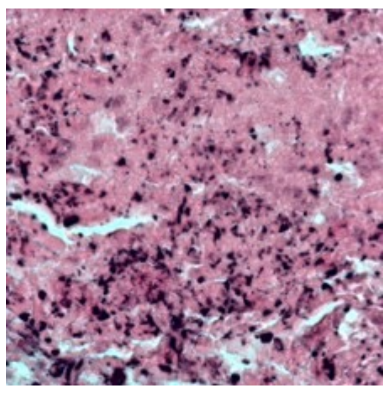

Dirty necrosis in colorectal cancer

HPC57

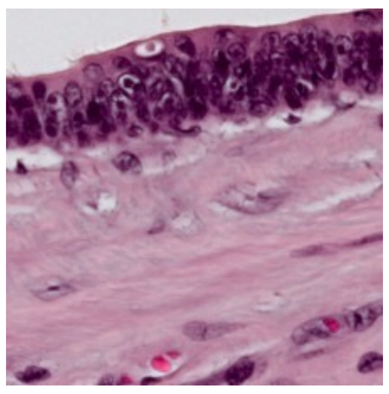

Heterogeneous cluster with sparse epithelial fragments, smooth muscle bundles, scattered inflammatory cells, and pale stroma with tissue artifacts, consistent with tumor edge or transitional zone between normal and neoplastic tissue.

HPC58

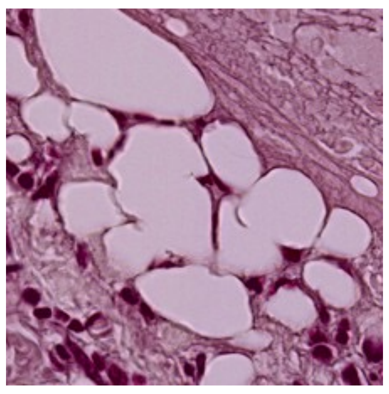

Normal colonic surface mucosa with goblet cell-rich crypts intimately admixed with underlying adipose tissue, consistent with mucosal-submucosal interface or serosal surface tiles capturing both epithelium and pericolorectal fat.

**Supplementary Figure 3 continued.**

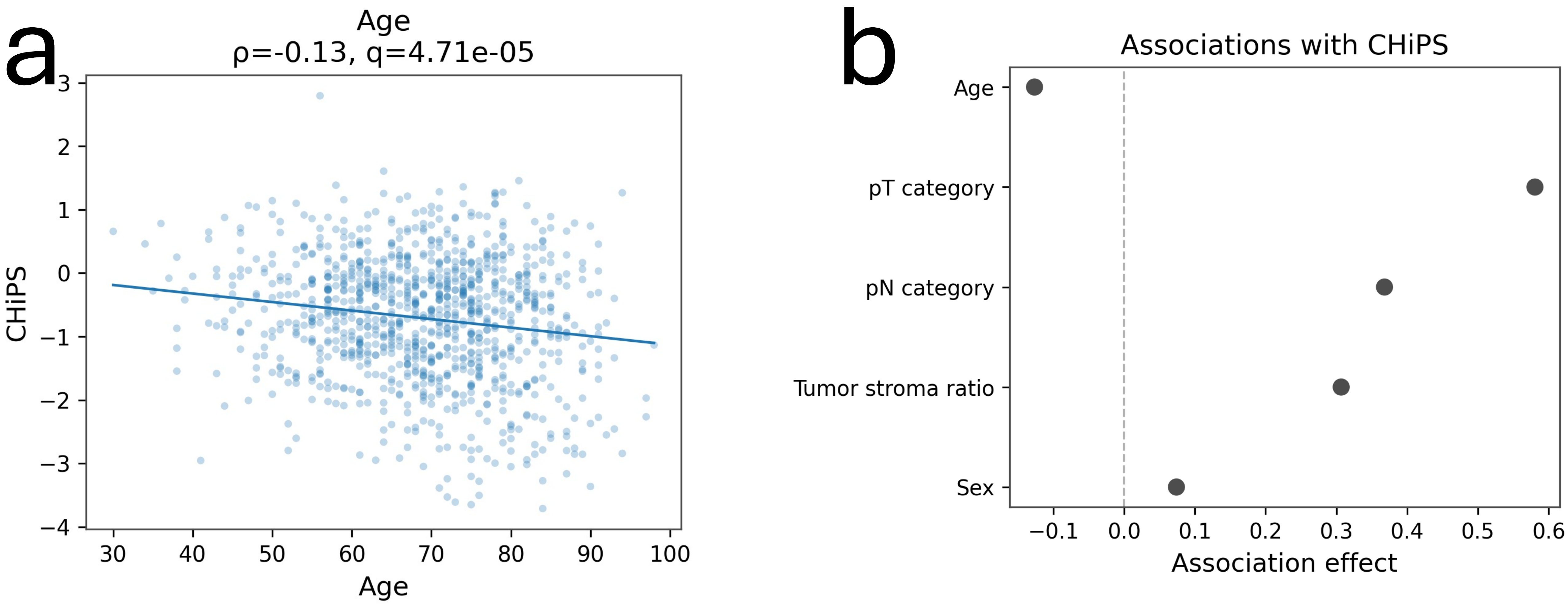


**Supplementary Figure 4 | Associations between CHiPS and clinicopathological variables.**

**a,** Association between CHiPS and age in the UNITED cohort. Each point represents one patient, and the fitted line shows the overall trend. CHiPS showed a weak negative correlation with age.

**b,** Effect sizes for associations between CHiPS and clinicopathological variables, including age, pathological T category, pathological N category, tumor stroma ratio, and sex. Pathological T category, pathological N category, and tumor stroma ratio showed stronger positive associations with CHiPS, whereas sex showed little association and age showed a weak negative association. CHiPS, Colon Histomorphology Prognostic Score.

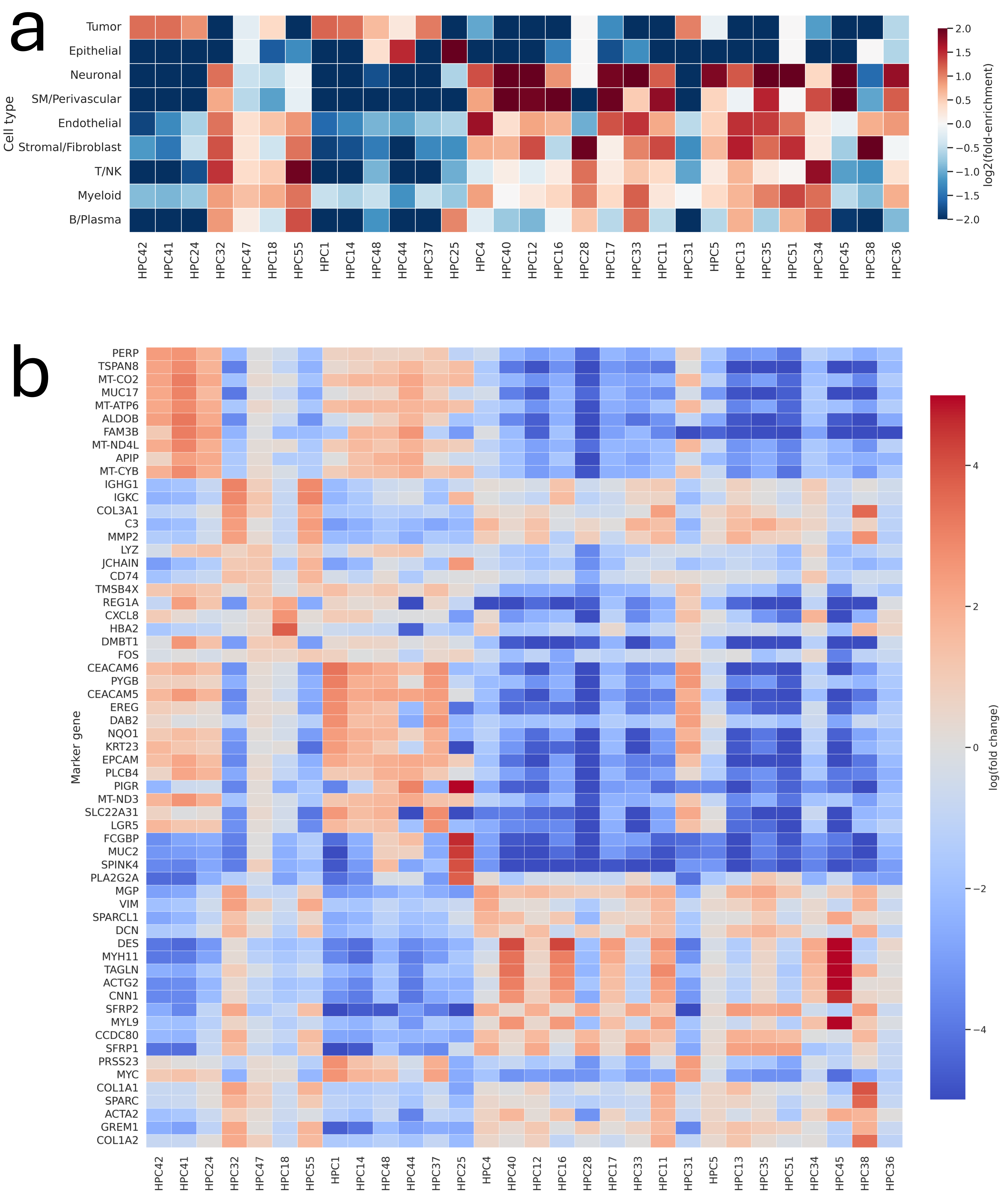
a
Tumor
Epithelial
Neuronal
SM/Perivascular
Endothelial
Stromal/Fibroblast
T/NK
Myeloid
B/Plasma
Cell type
log2(fold-enrichment)
HPC42 HPC41 HPC24 HPC32 HPC47 HPC18 HPC55 HPC1 HPC14 HPC48 HPC44 HPC37 HPC25 HPC4 HPC40 HPC12 HPC16 HPC28 HPC17 HPC33 HPC11 HPC31 HPC5 HPC13 HPC35 HPC51 HPC34 HPC45 HPC38 HPC36
b
Marker gene
PERP TSPAN8 MT-CO2 MUC17 MT-ATP6 ALDOB FAM3B MT-ND4L APIP MT-CYB IGHG1 IGKC COL3A1 C3 MMP2 LYZ JCHAIN CD74 TMSB4X REG1A CXCL8 HBA2 DMBT1 FOS CEACAM6 PYGB CEACAM5 EREG DAB2 NQO1 KRT23 EPCAM PLCB4 PIGR MT-ND3 SLC22A31 LGR5 FCGBP MUC2 SPINK4 PLA2G2A MGP VIM SPARCL1 DCN DES MYH11 TAGLN ACTG2 CNN1 SFRP2 MYL9 CCDC80 SFRP1 PRSS23 MYC COL1A1 SPARC ACTA2 GREM1 COL1A2
log(fold change)

**Supplementary Figure 5 | Cell-type enrichment and marker gene programs across CHiPS-associated histomorphological phenotype clusters.**

**a,** Broad cell-type enrichment across CHiPS-associated histomorphological phenotype clusters (HPCs), shown as log2 fold-enrichment relative to the global cell-type distribution across the analyzed Visium HD regions. Low-risk-associated epithelial/glandular HPCs showed enrichment for tumor and epithelial compartments, whereas high-risk-associated stromal/desmoplastic HPCs showed enrichment for stromal/fibroblast, smooth muscle/perivascular, endothelial, myeloid, or T/NK compartments.

**b,** Marker gene expression across CHiPS-associated HPCs. The top upregulated marker genes selected from the analyzed HPCs are shown as log fold change values across all displayed clusters, enabling comparison of marker specificity across epithelial, stromal, immune, and mesenchymal phenotypes. Epithelial/glandular clusters showed enrichment for tumor- and epithelial-associated markers, whereas stromal/desmoplastic clusters showed enrichment for fibroblastic, extracellular matrix, contractile, and immune-associated markers. CHiPS, Colon Histomorphology Prognostic Score; HPC, histomorphological phenotype cluster.

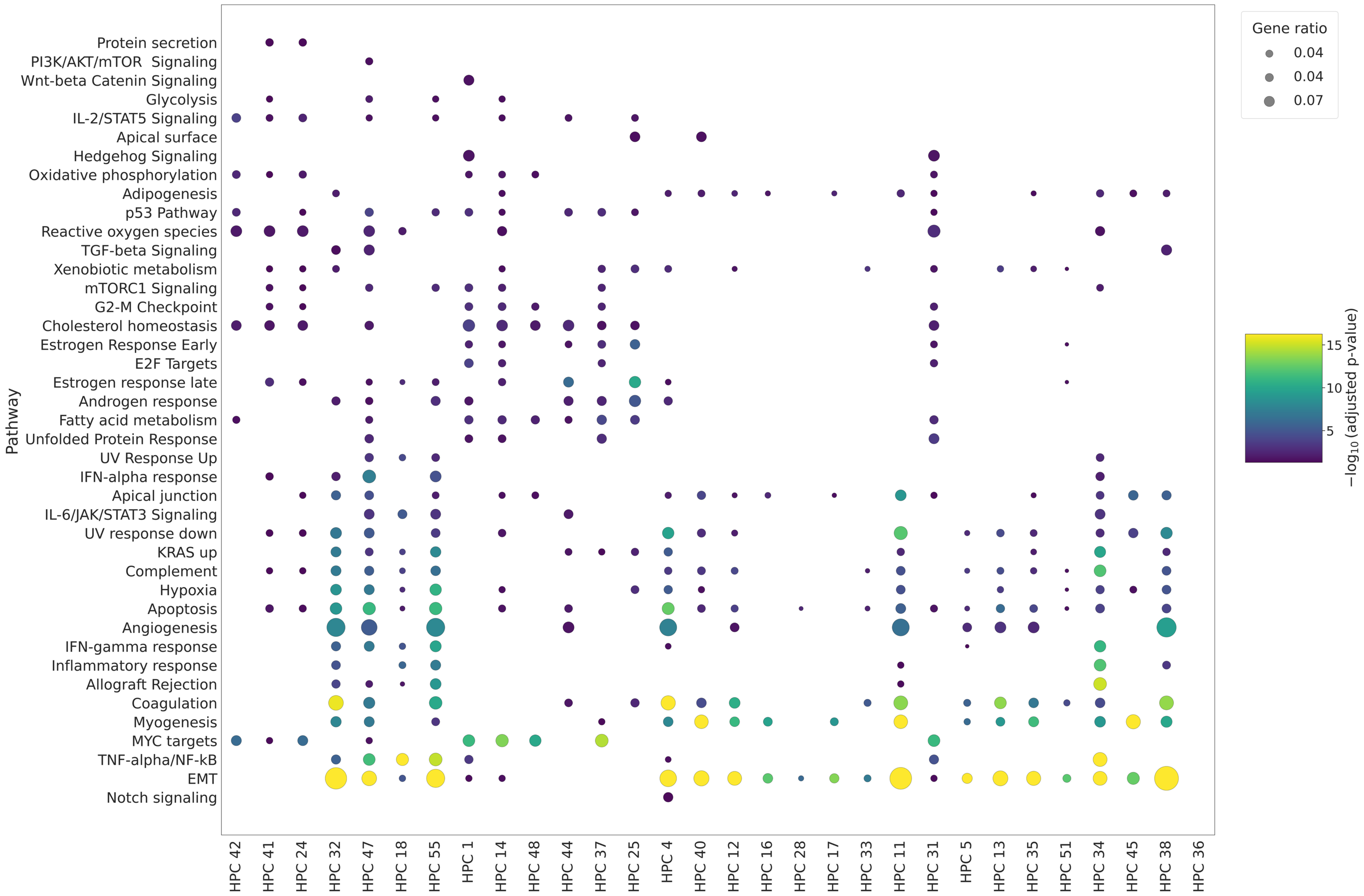


**Supplementary Figure 6 | Hallmark pathway enrichment across survival-associated histomorphological phenotype clusters.**

Hallmark pathway over-representation analysis (ORA) across CHiPS-associated histomorphological phenotype clusters (HPCs). ORA tests whether genes upregulated in each HPC are statistically over-represented in predefined Hallmark gene sets. Each column represents one HPC, and each row represents one Hallmark pathway. Dot size indicates the gene ratio, and dot color indicates enrichment significance as −log10 adjusted *P* value. Only significantly enriched pathways are shown. Low-risk-associated epithelial/glandular HPCs and high-risk-associated stromal/desmoplastic HPCs showed distinct pathway-level profiles, with high-risk-associated clusters showing recurrent enrichment for epithelial–mesenchymal transition, myogenesis, coagulation, hypoxia, inflammatory signaling, and extracellular matrix–related programs. This extended analysis supports the presence of molecularly distinct programs across the broader set of survival-associated HPCs.
